# NBLAST: Rapid, sensitive comparison of neuronal structure and construction of neuron family databases

**DOI:** 10.1101/006346

**Authors:** Marta Costa, James D. Manton, Aaron D. Ostrovsky, Steffen Prohaska, Gregory S. X. E. Jefferis

## Abstract

Neural circuit mapping is generating datasets of 10,000s of labeled neurons. New computational tools are needed to search and organize these data. We present NBLAST, a sensitive and rapid algorithm, for measuring pairwise neuronal similarity. NBLAST considers both position and local geometry, decomposing neurons into short segments; matched segments are scored using a probabilistic scoring matrix defined by statistics of matches and non-matches.

We validated NBLAST on a published dataset of 16,129 single *Drosophila* neurons. NBLAST can distinguish neuronal types down to the finest level (single identified neurons) without *a priori* information. Cluster analysis of extensively studied neuronal classes identified new types and unreported topographical features. Fully automated clustering organized the validation dataset into 1052 clusters, many of which map onto previously described neuronal types. NBLAST supports additional query types including searching neurons against transgene expression patterns. Finally we show that NBLAST is effective with data from other invertebrates and zebrafish.

## Introduction

Correlating the functional properties and behavioral relevance of neurons with their cell type is a basic activity in the study of neuronal circuits. While there is no universally accepted definition of neuron type, key descriptors include morphology, position within the nervous system, genetic markers, connectivity and intrinsic electrophysiological signatures (Migliore and Shepherd, 2005; Bota and Swanson, 2007; Rowe and Stone, 1976). Despite this ambiguity, neuron type remains a key abstraction helping to reveal organizational principles and enabling results to be compared and collated across research groups. There is increasing appreciation that highly quantitative approaches are critical to generate cell type catalogues in support of circuit research (Petilla Interneuron Nomenclature Group et al., 2008; Nelson et al., 2006; Kepecs and Fishell, 2014)(http://www.nih.gov/science/brain/11252013-Interim-Report-Final.pdf).

Since neuronal morphology and position strongly constrain connectivity, they have been mainstays of circuit studies for over a century. Classic morphological techniques include the Golgi method used by Cajal, microinjection, and intracellular fills during recording. Recently genetic approaches to sparse and combinatorial labeling have enabled increasingly large-scale characterization of single neuron morphology (Jefferis and Livet, 2012).

Classically, the position of neuronal somata or arbors was established by comparison with anatomical landmarks, revealed by a general counterstain; this is especially effective in brain regions with strong laminar organization e.g. the cerebellum (Cajal and Azoulay y, 1911), mammalian retina (e.g. Badea and Nathans, 2004; Kong et al., 2005; Sümbül et al., 2014) or fly optic lobe (Fischbach and Dittrich, 1989; Morante and Desplan, 2008). Recently, 3D light microscopy and image registration have enabled direct image fusion to generate digital 3D atlases of brain regions or whole brains (Jefferis et al., 2007; Lin et al., 2007; El Jundi et al., 2009; Rybak et al., 2010; Cachero et al., 2010; Yu et al., 2010b; Sunkin et al., 2013; Zingg et al., 2014; Oh et al., 2014). Such atlases can generate specific, testable hypotheses about circuit organization and connectivity at large scales. For example Chiang et al. (2011) combined genetic mosaic labeling and image registration to produce an atlas of over 16,000 single cell morphologies embedded within a standard *Drosophila* brain (http://flycircuit.tw).

Neuronal morphologies can be represented as directed graph structures embedded in 3D space; usually this is the (arbitrary) physical space of the imaging system, rather than a brain atlas. For this reason, databases such as NeuroMorpho.org (Parekh and Ascoli, 2013) contain > 27,000 neurons, but do not include precise positional information. Data on this scale present both an acute challenge, finding and organizing related neurons, but also an opportunity: quantitative morphology may help solve the problem of cell type. A key requirement is a tool enabling rapid and sensitive computation of neuronal similarity within and between datasets. This has clear analogies with bioinformatics: the explosion of biological sequence information from the late 80s motivated the development of sequence similarity tools such as BLAST (Altschul et al., 1990) enabling rapid database queries as well as hierarchically organized protein family databases.

Several strategies for measuring neuronal similarity exist with distinct target applications and different underlying data structures. Mayerich et al. (2012) applied general graph similarity metrics (reviewed by Conte et al., 2004) to compare neurons represented as fully-connected graphs with a ground truth reconstruction. Basu et al. (2011) decomposed neuronal trees into a family of unbranched paths proposing a geometric measure of similarity that could include positional information. Further simplifying the neuronal representation, Cardona et al. (2010) decomposed single unbranched neurites into sequences of vectors and used dynamic programming to find an optimal 3D alignment. Critically, they validated this approach on a database of a few hundred traced structures, achieving very high classification accuracy. However, this algorithm treats each unbranched neuronal segment as a separate alignment problem, so there is no natural way to handle trees with many such segments. Recently Wan et al. (2015) have developed an elegant approach that combines graph matching with 3D positional information for sensitive global alignment of reconstructed neurons, albeit at significant computational cost (minutes per pair of neurons).

The choice of data structure remains important: fully automatic 3D tracing of single neurons remains unsolved (Peng et al., 2015), while expression patterns containing multiple neurons cannot be represented as a single binary tree. We previously developed a pipeline that converts images consisting of 100s of neurons to point clouds with tangent vectors defining the local heading of the neurons; we used this as part of a supervised learning approach to the challenging problem of recognizing groups of lineage-related neurons (Masse et al., 2012). Combining this simple data representation with a very large single neuron dataset (Chiang et al., 2011) allowed us to validate a new algorithm, NBLAST, that is flexible, extremely sensitive and very fast (pairwise search times of 2 ms). Critically, the algorithm’s scoring parameters are defined statistically rather than by expert intuition, but generalize across neuronal classes.

We first describe the NBLAST algorithm, providing an open source implementation and a web query tool.We validate NBLAST for applications including neuron database search, unsupervised clustering and expression pattern search. NBLAST can identify well-studied neuronal types in *Drosophila* with sensitivity matching domain experts, in a fraction of the time. NBLAST can also identify new neuronal types and reveal undescribed features of topographic organization. Finally, we apply our method to 16,129 neurons from the FlyCircuit dataset, reducing this to a non-redundant set of 1,052 morphological clusters. Manual evaluation of a subset of clusters shows they closely match expert definition of cell types. These clusters, which we organize into an online supercluster hierarchy, represent a preliminary global cell type classification for the *Drosophila* brain.

## Results

### Algorithm

Our principal design goals were to develop a neuron similarity algorithm depending on both spatial location (within a brain or brain region) and branching pattern, and that was both extremely sensitive and very fast. We envisaged searching of large databases of neurons (10,000–100,000 neurons), clustering of neurons into families by calculating all-against-all similarity matrices, and efficient organization and navigation of datasets of this size. We eventually selected an approach based on direct pairwise of comparison of neurons pre-registered to a template brain and represented as vector clouds. Further details are provided in Supplemental Information.

The starting point for our algorithm is a representation in which neuron structures have been reduced to segments, represented as a location and an associated local tangent vector. This retains some local geometry but does not attempt to capture the topology of the neuron’s branching structure. We have found that a simplified representation of this sort can be constructed for image data that would not permit automated reconstructions. In order to prepare data of this sort in quantity, we developed an image processing pipeline summarized in Figure 1A and detailed in Experimental Procedures. Briefly, brain images from the FlyCircuit dataset (Chiang et al., 2011) were subjected to non-rigid image registration (Jefferis et al., 2007) against a newly constructed intersex template brain. Neuron images were thresholded and subjected to a 3D skeletonization procedure (Lee et al., 1994) implemented in Fiji (Schindelin et al., 2012). These thresholded images were then converted to the point and tangent vector representation (Masse et al., 2012) using our R package [nat](http://cran.r-project.org/web/packages/nat/index.html) (Jefferis and Manton, 2014); the tangent vector (i.e. the local heading) of the neuron at each point was computed as the first eigenvector of a singular value decomposition (SVD) of the point and its 5 nearest neighbors.

**Figure 1:**
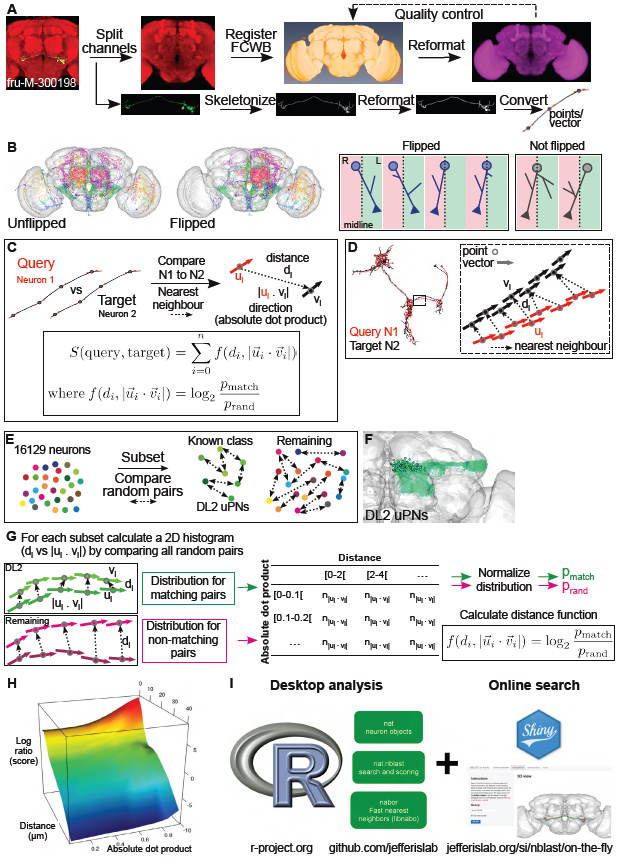
Image preprocessing, registration and similarity score (NBLAST) algorithm. **(A)** Flowchart describing the image preprocessing and registration procedure. Fly-Circuit images were split into 3 channels. The Dlg-stained brain (discs large 1) images were registered against the FCWB template. Successful registrations were applied to the skeletonized neuron images. Neuron skeletons were converted into points and vectors. **(B)** Neurons on the right side of the brain were flipped onto the left. Brain plots showing 50 random neurons before and after flipping. On the right, cases for which the neuron flipping was assessed manually. **(C)** NBLAST algorithm. The similarity of two neurons (query and target), is given by a function of the distance and absolute dot product between the nearest neighbor points of the query/target pair. This distance function reflects the probability of a match between a pair of points (p**_match_**), relative to any two random points (p**_rand_**). **(D)** Diagram illustrating how nearest neighbor points are calculated. For a query (N1)/target (N2) pair, each point of N1 (u_i_) is paired to the N2 point (v_i_) that minimizes the distance (d_i_) between the points. **(E)** Calculating the distance function. Random pairs of neurons within two groups, DL2 uPNs and all remaining neurons, were compared. **(F)** Brain plot of DL2 uPNs. **(G)** Calculation of the distribution for matching and non-matching pairs of segments. For all segment pairs of all neuron pairs of each group, the distance and absolute dot product were plotted in a distance histogram. The distribution probability for matching (p**_match_**) or non-matching pairs (p**_rand_**) was calculated by normalizing the distance histogram to 1. **(H)** Plot showing that the similarity score depends on the spatial location of the points (distance between points) and the direction of the vectors (absolute dot product). **(I)** Summary diagram of the NBLAST implementation, for desktop and online uses.

After pre-processing, 3D data could be visualized and analyzed in R using nat (Figure 1B). Neurons were represented by median 1,070 points/vectors; the 16,129 neurons occupied 1.8 GB, fitting comfortably into a laptop’s main memory. Since the fly brain is almost completely symmetric, but neurons were labeled randomly in both hemispheres, we mapped all neurons to the left hemisphere (defined primarily by cell body location, see Experimental Procedures and Figure 1B) using a nonrigid mirroring procedure (Manton et al., 2014).

With a database of aligned neurons in an appropriate representation, we were then able to calculate NBLAST pairwise similarity scores. One neuron is designated the query and the other the target. For each query segment (defined by a midpoint and tangent vector) the nearest neighbor (using straightforward Euclidean distance) is identified in the target neuron (Figure 1C–D). A score for the segment pair is calculated as a function of two measurements: *d*, the distance between the matched segments (indexed by *i*), and 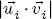, the absolute dot product of the two tangent vectors; the absolute dot product is used because the orientation of the tangent vectors has no meaning in our data representation (Figure 1C). The scores are then summed over each segment pair to give a raw score, *S*:

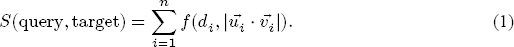

The next question is what is an appropriate function 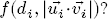 We developed an approach inspired by the scoring system of the BLAST algorithm (Altschul et al, 1990). For each segment pair we defined the score as the log probability ratio:

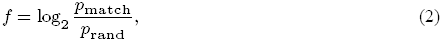

i.e. the probability that the segment pair was derived from a pair of neurons of the same type, versus a pair of unrelated neurons. We could then define *p*_match_ empirically by finding the joint distribution of *d* and 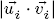 for pairs of neurons of the same type (Figure 1E–G). For our default scoring matrix, we used a set of 150 olfactory projection neurons innervating the same glomerulus, unambiguously the same neuronal type (Figure 1F). *p*_rand_ was calculated simply by drawing 5,000 random pairs of neurons from the database, assuming that the large majority of such pairs are unrelated neurons. Joint distributions were calculated using 10 bins for the absolute dot product and 21 bins for the distance to give two 21 row × 10 column matrices. The 2D histograms were then normalized to convert them to probabilities and the log ratio defined the final scoring matrix (Figure 1G). Plotting the scoring matrix emphasizes the strong distance dependence of the score but also shows that for segment pairs closer than ~10 μm, the logarithm of the odds score increases markedly as the absolute dot product moves from 0 to1 (Figure 1H).

We implemented the NBLAST algorithm as an R package (nat.nblast) building on a high-performance k-nearest neighbor library ([nabor](http://cran.r-project.org/package=nabor)), that immediately enables pairwise queries, searches of a single query neuron against a database of target neurons (Figure 2) and all-by-all searches (Figure 1I). Runtimes on a single core laptop computer were 2 ms per comparison or 30 s for all 16,129 neurons. In order to enable interactive neuron clustering, we also pre-computed an all-by-all similarity matrix for all 16,129 neurons (2.6 × 10^8^ scores, 1.0 GB). We also developed a simple web application (linked from jefferislab.org/si/nblast) to allow online queries for this test dataset (Figure 1I).

**Figure 2:**
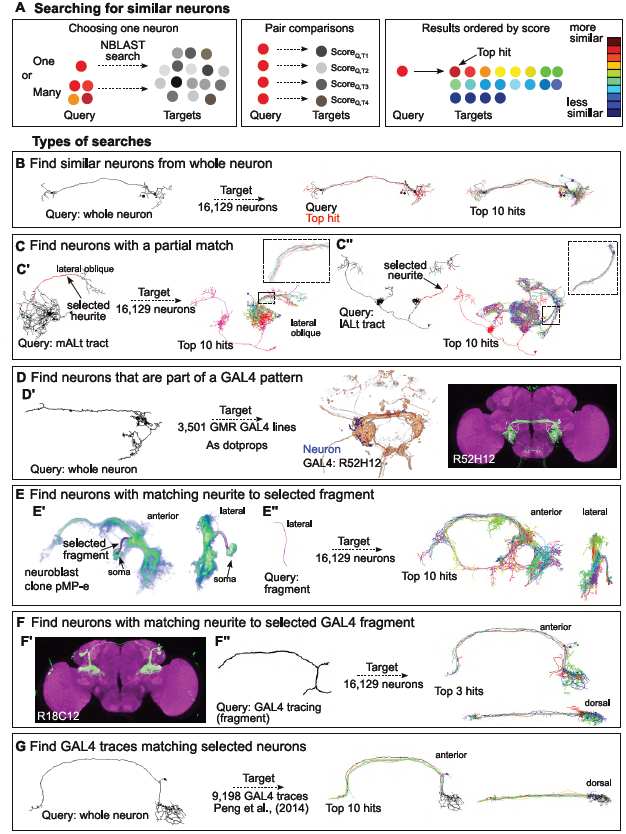
NBLAST allows different types of searches. **(A)** Searching for neurons with NBLAST. Pair comparisons between the query and target neurons in the dataset return a similarity score, allowing the results to be ordered. **(B)** NBLAST search using a whole neuron as a query against the FlyCircuit dataset. The query neuron (fru-M-400121), top hit and top 10 hits in anterior view. **(C)** NBLAST search using a neuron fragment as a query against the FlyCircuit dataset. The query neuron and top 10 hits are shown. **(C’)** Search with the mALT tract from an olfactory PN (Cha-F-000239). Lateral oblique view is shown, the inset shows the mALT tract of the top 10 hits. **(C”)** Search with the lALT tract from an olfactory PN (Gad1-F-200095). Anterior view is shown, the inset shows the lALT tract of the top 10 hits. **(D)** NBLAST search against FlyLight GAL4 lines using a query neuron. **(D’)** From left to right: query neuron (Trh-M-300069), volume rendering of query and best GAL4 hit (R52H12), maximum Z projection of line R52H12. **(E)** NBLAST search for neurons with matching neurites to a fragment from a *fruitless* neuroblast clone (pMP-e). The target is the FlyCircuit dataset. **(E’)** Volume rendering of pMP-e clone with the selected fragment – the characteristic stalk of P1 neurons. An anterior and lateral view are shown. **(E”)** Query fragment in lateral view. Top 10 hits in anterior and lateral view. **(F)** NBLAST search for neurons with matching neurites to a fragment traced from a GAL4 image (R18C12) ((Jenett et al., 2012)). The target is the FlyCircuit dataset. **(F’)** Maximum Z projection of line R18C12. **(F”)** The fragment used as query (traced in Vaa3D) in anterior view. Top 3 hits in anterior and dorsal view. **(G)** NBLAST search for GAL4 traces (Peng et al., 2014) matching a selected FlyCircuit neuron (VGlut-F-500818). The top 10 trace hits are shown in anterior and dorsal view.

### NBLAST finds whole or partial matches for diverse query objects

The NBLAST algorithm is flexible, identifying both global and partial matches for multiple classes of queries (Figure 2). The only requirements are that these objects (or fragments) must be registered against a template brain and can be converted to a point and vector representation.

As a first example we query a (whole) FlyCircuit neuron against the 16,129 FlyCircuit neurons. The top hits are very similar neurons with small differences in length and neurite position (Figure 2B). A second example uses part of the axon; top hits all follow the same axon tract, although their variable axonal and dendritic arbors indicate that they are distinct neuron types (Figure 2C). In a third example we query against 3501 FlyLight GAL4 lines (Jenett et al., 2012), finding lines that contain the query neuron (Figure 2D).

User tracings can also be used as queries. We traced the characteristic bundle of 20-30 primary neurites of the *fruitless* neuroblast clone pMP-e (which generates male-specific P1 neurons, (Kimura et al., 2008; Cachero et al., 2010)). Our query returned many single P1 neurons from the FlyCircuit database (Figure 2E) (for more details see Figure 7). A similar approach can be used to identify candidate neuronal types labeled by genetic driver lines where the detailed morphology of individual neurons cannot be ascertained. In this example we traced the main neurites of a cell cluster in a GAL4 line(Jenett et al., 2012) (Figure 2F) and used that trace as the query. NBLAST identified three very similar FlyCircuit neurons, which completely overlapped with the expression of GAL4 line. These three neurons appear to be different subtypes, each varying in their terminal arborizations. Conversely, we used one tracing from a published projectome dataset containing > 9000 neurite fibers (Peng et al., 2014) to find similar FlyCircuit neurons (Figure 2G).

### NBLAST scores are sensitive and biologically meaningful

A good similarity algorithm should be sensitive enough to reveal identical neurons with certainty, while having the specificity to ensure that all high scoring results are relevant. We used the full FlyCircuit dataset to validate NBLAST performance.

Our first example uses an auditory interneuron, [fru-M-300198](http://flycircuit.tw/modules.php?name=clearpage&op=detail_table&idid=516) as query (Figure 3A–C). Ordering search results by NBLAST score, the first returned object is the query neuron itself (since it is present in the database), followed by the top hit ([fru-M-300174](http://flycircuit.tw/modules.php?name=clearpage&op=detail_table&idid=427)) which completely overlaps with the query (Figure 3A’). A histogram of NBLAST scores, showed that the top hit was clearly an outlier: scoring 96.1 % compared to the self-match score of the query neuron (Figure 3C). Further investigation revealed that these “identical twins”, both derived from the same raw confocal image. The next 8 hits are also very similar to the query but are clearly distinct specimens, having small differences in position, length and neurite branching that are typical of sister neurons of the same type (Figure 3A”).

**Figure 3:**
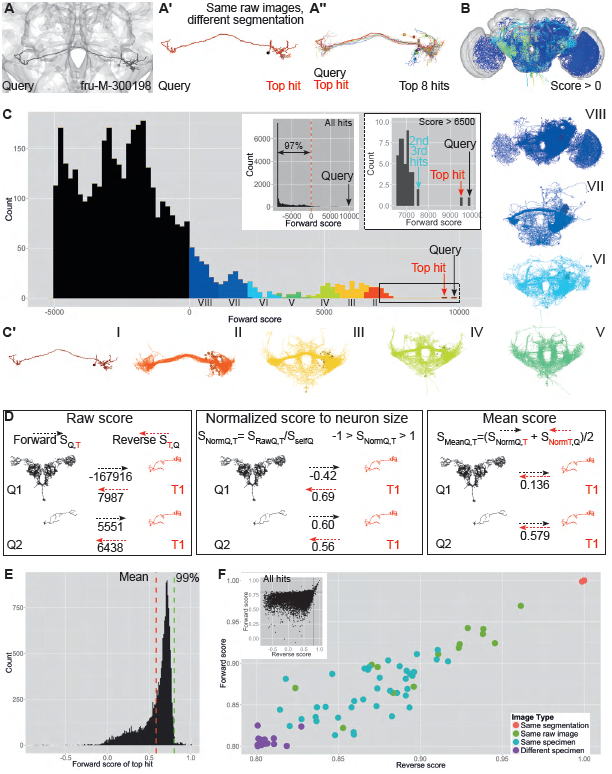
NBLAST scores are accurate and meaningful. **(A)** NBLAST search with fru-M-300198 (black). **(A’)** Neuron plot of the query (black) and top hit (red). The top hit corresponds to a different segmentation of the query, from the same raw image. **(A”)** Neuron plot showing the top 8 hits. There are differences in neurite branching, length and position. **(B)** All hits with a forward score over 0, colored by score, asshown inC. **(C)** Histogram of scores for a forward search with fru-M-300198 as query. Only hits with scores over -5,000 are shown. The left inset shows the histogram of scores for all search hits. The right inset shows a zoomed view of the top hits (score > 6,500). For more examples see **S1**. **(C’)** Neuron plots corresponding to the score bins in C. **(D)** Comparison of the raw, normalized and mean score, for two pairs of neurons: one of unequal (Q1, T1) and one of similar size (Q2, T1). **(E)** Histogram of the normalized score for the top hit for each neuron in the whole dataset. The mean and 99th percentile are shown as a dashed red and green lines, respectively. **(F)** Plot of reverse and forward normalized scores for 72 pairs of neurons for which both the forward and reverse scores are higher than 0.8. These pairs were classified into four categories: images correspond to a segmented image that is duplicated (‘Same segmentation’); images correspond to different neuron segmentations from the same raw image (‘Same raw image’); images correspond to two different segmented images from the same brain (‘Same specimen’); images correspond to segmented images of the same or similar neurons in different brains (‘Different specimen’). The inset plot shows the normalized reverse and forward scores for all top hits. The threshold of 0.8 is indicated by two black lines.

The score histogram shows that only a minority of hits (3 %) have a score above 0 (Figure 3B–C). A score of 0 represents a natural cutoff for NBLAST, since it means that, on average, segments from this neuron have a similarity level that is equally likely to have arisen from a random pair of neurons in the database as a pair of neurons of the same type. We divided the neurons with score > 0 into 8 groups with decreasing similarity scores (Figure 3C’). Only the highest scoring real hits (group II) appear of exactly the same type, although lower scoring groups contain neurons that would be ranked as very similar.

Although raw NBLAST scores correctly identify similar neurons, they are not comparable from one query neuron to the next: the score depends on neuron size and segment number. This confounds search results for neurons of very different sizes or when the identity of query and target neurons is reversed. For example, a search with a large neuron as query and a smaller one as target (pair 1) will have a very low **forward** score, because the large neuron has many segments that are unmatched, but a high **reverse** score, since most of target will match part of the query (Figure 3D). One approach to correct for this is to normalize the scores by the size of the query neuron. Although **normalized** scores are comparable, unequal forward and reverse scores between large and small neurons remain an issue. One simple strategy is to calculate the mean of the forward and reverse scores (**mean score**). Two neurons of similar size have a higher mean score than two neurons of unequal size (Figure 3D). Repeating the analysis of Figure 3C–C’ using mean scores (Figure S2) eliminated some matches due to unequal size that could be considered false positives.

During our analysis, we sporadically noticed cases where two database images were derived from the same physical specimen (Figure S1). We tested if NBLAST could systematically reveal such instances. We collected the top hit for each neuron and analyzed the distribution of forward (Figure 3E) and reverse scores (data not shown). A small tail (~ 1 % of all top hits) have anomalously high scores (over 0.8). Given this distribution, we examined neuron pairs with forward and reverse scores >0.8. We classified these 72 pairs into 4 different groups. From highest to lowest predicted similarity, the groups are: same segmentation, a segmented image of a neuron has been duplicated (Figure S1A); same raw image, corresponding to a different segmentation of the same neuron (Figure 3B’); same specimen, when two images are from the same brain but not from the same confocal image (Figure S1B); and different specimen, when two neurons are actually from different brains, (but of the same neuron type). The distribution of NBLAST scores for these four categories matches the predicted hierarchy of similarity (Figure 3F). These results underline the high sensitivity of the NBLAST algorithm to small differences between neurons.

Taken together these results validate NBLAST as a sensitive and specific tool for finding similar neurons.

### NBLAST scores can distinguish Kenyon cell classes

We wished to investigate whether NBLAST scores can be used to cluster neurons, potentially revealing functional classes. We decided to begin our investigation with Kenyon cells (KCs), the intrinsic neurons of the mushroom body neuropil and an intensively studied population given their key role in memory formation and retrieval (reviewed in Kahsai and Zars, 2011).

There are around 2,000 KCs in each mushroom body Aso et al. 2009, and their axons form the medial lobe, consisting of the γ, β′ and β lobes, and the vertical lobe, consisting of the α and α′ lobes. There dendrites form the calyx around which cell bodies are positioned; the axon peduncle joins the calyx to the lobes (Figure 4A). Three main classes of KCs and a few subclasses of neurons are recognized: γ neurons are the first born and innervate only the γ lobe; α′/β′ neurons are generated next and project to the α′ and β′ lobes; α/β neurons are born last, and project to the α and β lobes. Four neuroblasts each generate the whole repertoire of KC types (Lee et al., 1999).

**Figure 4:**
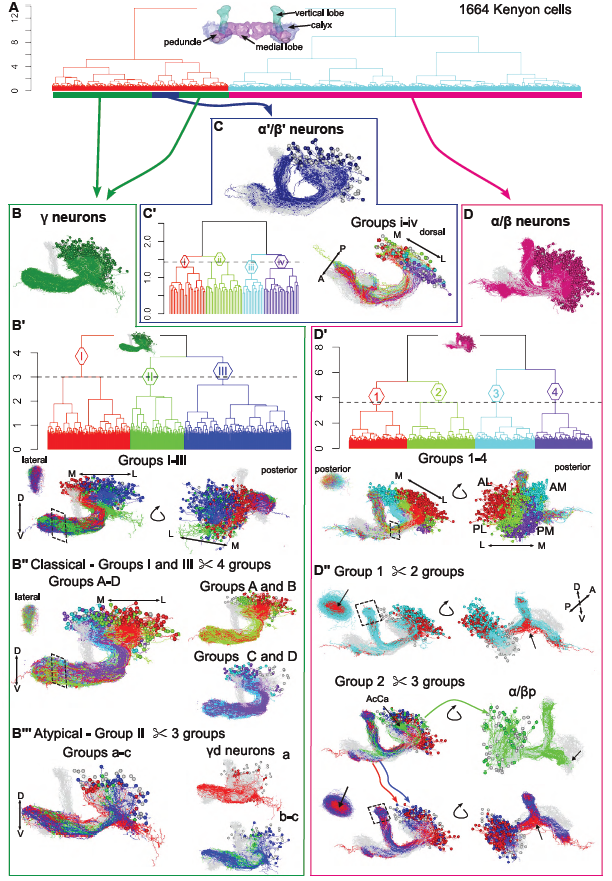
NBLAST search and classification of hits reveals Kenyon cell subtypes. **(A)** Hierarchical clustering (HC) of Kenyon cells (n=1664). Bars below the dendrogram indicate the γ (in green), α′/β′ (in blue) and α/β neurons (in magenta), h=8.9. Inset shows the mushroom body neuropil. **(B)** Neuron plot of γ neurons. **(B’)** HC of γ neurons (I-III), h=3. Neuron plots of groups I to III. A lateral oblique and a posterior view of the neurons and a lateral view of a horizontal lobe slice are shown. **(B”)** HC of the classic γ neurons, corresponding to groups I and III in B’, divided into four groups (A–D). Neuron plots of groups A–D, A–B and C–D. **(B”’)** HC of atypical γ neurons corresponding to group II in B’, divided into three groups (a–c). Neuron plots of groups a–c, a, and b–c. Group a corresponds to subtype γd neurons. **(C)** Neuron plot of α′/β′ neurons. **(C’)** HC of α′/β′ neurons, divided into four groups (i–iv), h=1.43. **(D)** Neuron plot of α/β neurons. **(D’)** HC of α/β neurons, divided into four groups (1–4), h=3.64. Neuron plots of groups 1 to 4, which correspond to the neuroblast clones (AM, AL, PM, PL). Lateral oblique, posterior view and posterior view of a peduncle slice of these groups are shown. **(D”)** HC of groups 1 and 2. Lateral oblique, posterior oblique and a dorsal view of a peduncle slice views are shown. HC of group 1 divided into 2 subgroup, separating the neurons into surface (cyan) and core (red). Similar analysis to groups 3 and 4 is shown in Figure S3A. HC of group 2 divided into 3 subgroups. The red and blue subgroups match the core and surface neurons, respectively; the green subgroup corresponds to the α/β posterior subtype (α/βp). AcCa: accessory calyx. Neurons in grey: Kenyon cell exemplars.

We started with a dataset of 1,664 KCs, representing 10.3 % of the FlyCircuit dataset (see Supplemental Results for selection protocol) and calcaulted raw NBLAST scores of each KC against all others. An iterative hierarchical clustering approach allowed us to identify the main KC types, followed by detailed analysis for that distinguished several subtypes for each type.

For γ neurons (Figure 4B’), we identified 2 subsets, one corresponding to the classical morphology (Figure 4B”) (groups I and III) and another to recently described atypical neurons (γ dorsal neurons, group II) (Aso et al., 2014). Analysis of the classical γ neurons revealed that there were differences between the neurons in their medial to lateral position in the calyx (groups A-D). These differences correlated to a certain degree with differences in the dorsal/ventral position of the projections in the γ lobe, with the most medial, being also the most dorsal (Figure 4B”). These observations suggest that the relative position of the projections of classical γ neurons is maintained at the calyx and γ lobe. We experimented with clustering the classic γ neurons based only on the scores of the segments in the peduncle. The overall organization almost fully recapitulated the positioning of the neurites in the whole neuron analysis (see Figure S3 and). Thus, the stereotypical organization of the classical γ neurons is maintained throughout the neuropil.

The atypical γ neurons extended neurites posteriolaterally in the calyx and projected to the most dorsal region of the γ lobe (Figure 4B”’). We isolated a previously identified subtype – γd neurons (group a) (Aso et al., 2009) – that innervates the ventral accessory calyx (Aso et al., 2014). In addition, we identified previously uncharacterized types (Groups b-c).

Analysis of α′/β′ neurons highlighted the characterized subtypes of these neurons (Figure 4C–C’) which differ in their anterior/posterior position in the peduncle and β′ lobe (Tanaka et al., 2008; Aso et al., 2014). Although we could not identify any unambiguous subtypes of α′/β′ neurons, there were clear trends, with a subset of neurons (groups ii, iii) having more anterior neurites in both the peduncle and β′ lobe.

The largest KC subset corresponds to α/β neurons (Figure 4D). We identified neurons from each of the four neuroblast lineages (Figure 4D’) (Zhu et al., 2003) and for each of these, we distinguished morphological subtypes that correlate to their birth time (Figure 4D”). There was a clear distinction between the late born core (α/β core, α/β-c), on the inside stratum of the α lobe, and early born peripheral neurons (α/β surface, α/β-s), on the outside stratum of the α lobe. We also identified the earliest born α/β neurons – α/β posterior or pioneer (α/βp) – that innervate the accessory calyx and run along the surface of the posterior peduncle into the β lobe but stop before reaching the medial tip (Tanaka et al., 2008). A new clustering based on peduncle position of the neuron segments did not recapitulate the relative positions of the calyx neurites for each of the neuroblast clones observed in the whole neuron analysis suggesting that the relative position of the α/β neurons in the peduncle does not completely reflect their stereotypical organization in the calyx (see Figure S3 and Kenyon cell analysis).

In summary, the hierarchical clustering of KCs using the raw NBLAST scores resolved the neurons into the previously described KC types and some of the subtypes, and isolated uncharacterized subtypes in an extensively studied cell population. In addition, it revealed organizational principles that have been previously described (Tanaka et al., 2008). These observation support our claim that the NBLAST scores are a good metric when searching for similar neurons and organizing large datasets of related cells.

### NBLAST identifies classic cell types at the finest level: olfactory projection neurons

We have shown that clustering NBLAST scores can identify the major classes and subtypes of Kenyon cells. However it is rather unclear what corresponds to an identified cell type, which we take to be the finest neuronal classification in the brain. We therefore analyzed a different neuron family, the olfactory projection neurons (PNs), which represent one of the best defined cell types in the fly brain.

PNs transmit information between antennal lobe glomeruli, which receive olfactory input, and higher brain centers, including the mushroom body and the lateral horn (Masse et al., 2009). Uniglomerular PNs (uPNs), whose dendrites innervate just one glomerulus, are highly stereotyped in both morphology and developmental origin. They are unambiguously classified into individual types based on the glomerulus they innervate and the axon tract they follow; these features show fixed relationships with their axonal branching patterns in higher centers and their parental neuroblast (Marin et al., 2002; Jefferis et al., 2001; Wong et al., 2002; Jefferis et al., 2007; Yu et al., 2010a; Tanaka et al., 2012).

We manually classified the 400 FlyCircuit uPNs by glomerulus in an iterative process that took several days (for details see). We found a very large number of DL2 uPNs (145 DL2d and 37 DL2v), in a total of 397 classified neurons. Nevertheless, our final set of uPNs broadly represents the total variability of described classes and contains neurons innervating 35 out of 56 different glomeruli (Tanaka et al., 2012), as well as examples of the three main lineage clones and tracts.

We computed mean NBLAST scores for each uPN against the remaining 16,128 neurons and checked whether the top hit was exactly the same type of uPN, another uPN or a match to another class of neuron (Figure 5A). We restricted our analysis to types with at least two examples in the dataset and to unique pairs (if PN A and PN B were top hits for each other, we only counted them once) (n=327). There were only 8 cases in which the top hit did not match the query’s class. Of these, four had matches to a uPN innervating a neighboring glomerulus with identical axon projections (DL2d vs DL2v, VM5d vs VM5v) that are challenging even for experts to distinguish. There were a further four matches to multi-glomerular PNs that innervated the same glomerulus as well as some others. This exercise encapsulates a very simple form of supervised learning (k-nearest neighbor with k=1 and leave one out cross-validation) and shows that NBLAST scores are a suitable metric for this form of machine learning with an error rate of 2.4% for 35 classes; it also noteworthy that this was in the face of a huge amount of distracting information since uniglomerular PNs represented on only 2.47% of the 16128 test neurons.

**Figure 5:**
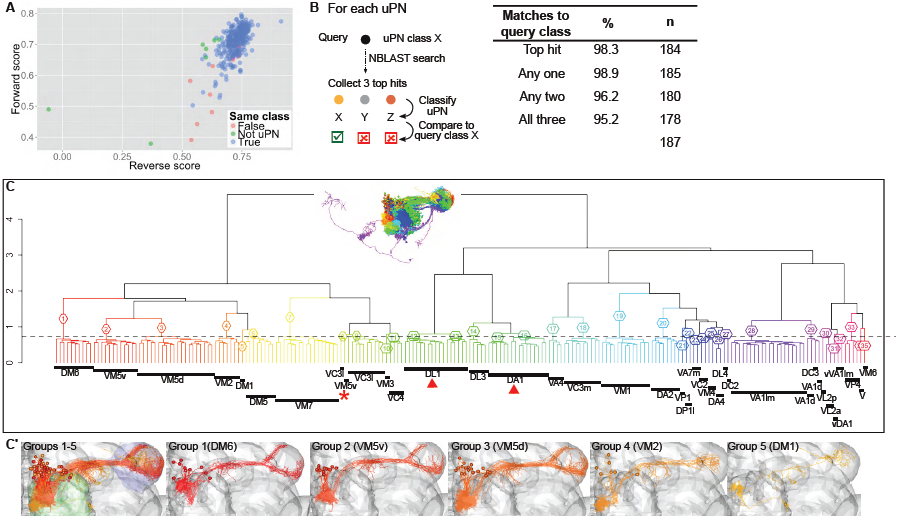
NBLAST search and classification of hits reveals uniglomerular olfactory projection neuronal types. **(A)** Plot of the reverse and forward normalized scores for the top hit in an NBLAST search using the uniglomerular olfactory projection neurons (uPNs) as queries (n=327). For each query neuron, we identified cases for which both the top hit and query were of the same class (True) (n=319); the top hit is a uPN but does not match the class of the query (False) (n=4), or the top hit is not a uPN (Not uPN) (n=4). **(B)** Percentage of neuron type matches in the top hit and top 3 hits for each uPN. The top three hits for each uPN (mean score) were collected and the neuron type of each hit and query was compared.**(C)** Hierarchical clustering of uPNs (non-DL2s) (n=214) divided into 35 groups (1–35), h=0.725. Dendrogram showing the innervated glomerulus for each neuron. Inset shows the uPNs colored by dendrogram group. Below the leaves, the number of neurons that innervate each glomerulus is indicated by the black rectangles. Neurons that innervate DA1 and VA1lm glomeruli but originate from the ventral lineage instead of the lateral or anterodorsal, respectively, are indicated as vVA1lm and vDA1. The dendrogram groups correspond to single and unique neuron types except for DL1 and DA1 neurons which are split into 2 groups (12–13, 15–16, respectively) (red arrowhead) and the outlier neuron VM5v in group 9 (red asterisk). **(C)** Neuron plots corresponding to dendrogram groups 1–5 and to each of these individual groups. The antennal lobe is in green, the lateral horn in purple.

We also compared how the top 3 hits matched the query type (Figure 5B). For uPN types with more than three examples (non DL2, n=187), we collected the top three NBLAST hits for each of these neurons. We achieved very high matching rates: in 98.9 % of cases (i.e. all except two) at least one of the top hits matched the query type, and all three hits matched the query type in 95.2% of cases.

Given the very high prediction accuracy, we wondered if an unsupervised clustering based on NBLAST mean scores would group uPNs by type. To test this, we clustered uPNs (non DL2, n=214) and divided the dendrogram at a height of 0.725: as this level most groups corresponded to single and unique neuron types. For types with more than one representative neuron, all neurons co-clustered, with three exceptions (Figure 5C). In one case there was substantial mis-registration of the antennal lobe overlooked by our image registration quality control; in the other 2 cases, neurons of the same type were split into two adjacent clusters. The cluster organization also reflects higher level features such as the axon tract / neuroblast of origin (Figure 5C’). Thus, unsupervised clustering of uPNs based on NBLAST scores gives an almost perfect neuronal classification: our two expert annotators took three iterative rounds of consensus-driven manual annotation to better this error rate of 1.4%.

In conclusion, these results demonstrate that morphological comparison by NBLAST is powerful enough to resolve differences at the finest level of neuronal classification. Furthermore, they suggest that unsupervised clustering by NBLAST scores could help reveal new neuronal types.

### NBLAST can be used to define new cell types

#### Visual projection neurons

Visual projection neurons (VPNs) relay information between optic lobe and the central brain. They are a morphologically diverse group with 44 types already described (Otsuna and Ito, 2006). We explored whether clustering of these neurons based on NBLAST scores would find previously reported neuron classes and identify new ones.

We isolated a set of VPNs including 1,793 unilateral VPNs, 72 bilateral VPNs and 2,892 intrinsic optic lobe neurons. Hierarchical clustering of the unilateral VPNs (uVPNs) resulted in a dendrogram, which we divided into 21 groups (I-XXI), in order to isolate one or a few cell types by group based both directly on morphological stereotypy and on previous literature (Otsuna and Ito, 2006) (Figure 6A–A’ and Figure S4A–A’). We further investigated these groups to determine if central brain innervation was a major differentiating characteristic between classes.

**Figure 6:**
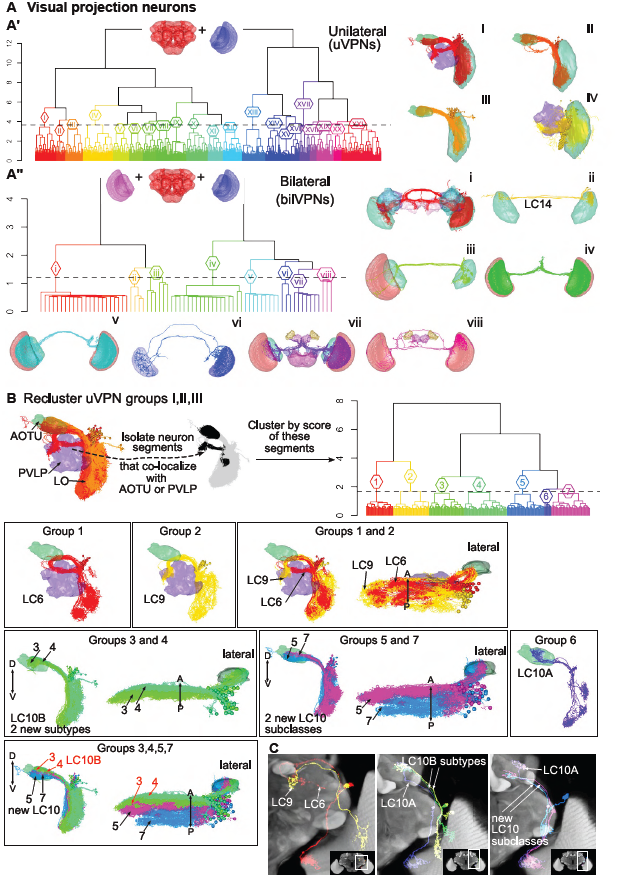
NBLAST search and classification of hits uncovers visual projection neuronal types. **(A)** Clustering analysis of unilateral (uVPNs) and bilateral visual projection neurons (bilVPNs). Inset on the dendrogram shows the neuropils considered for the overlap. To the right, neuron plots of the indicated groups, showing the neuropils with the most overlap. Other neuron plots are shown in Figure S4. **(A’)** Hierarchical clustering (HC) of uVPNs, divided into 21 groups (I–XXI), h=3.65. **(A”)** HC of bilVPNs, divided into 8 groups (i–viii), h=1.22. Group ii corresponds to the LC14 neuron type. **(B)** Reclustering of uVPN groups I, II and III from A’. The neuron segments that co-localize with either the anterior optic tubercle (AOTU) or posterior ventrolateral protocerebrum (PVLP) were isolated, followed by HC of the neurons based on the NBLAST score of these neuron segments. The dendrogram was divided into seven groups (1–7), h=1.69. Below, neuron plots matching the dendrogram groups to known uVPN types neuron types. Group 1: LC6 neurons; group 2: LC9. Groups 3 and 4: two new subtypes of LC10B. Groups 5 and 7: possible new subclasses of LC10. Group 6: LC10A neurons. **(C)** Overlay of Z projections of registered image stacks of example neurons from the types identified in B on a partial Z projection of the template brain (a different one for each panel). The white rectangle on the inset shows the location of the zoomed in area. LC: lobula columnar neuron.

**Lobula-, AOTU- and PVLP-innervating uVPNs** We took neuron skeletons from groups I–III uVPN and isolated the axon arbors innervating the anterior optic tubercle (AOTU) and posterior ventrolateral protocerebrum (PVLP) (Figure 6B). A new clustering based on the scores of these partial skeletons, allowed us to identify 7 different groups (1–7). A clear distinction between neurons that innervated the PVLP (groups 1, 2) and those that extensively innervated the AOTU (groups 3–6) was evident. Our analysis divides the LC10 uVPN class into 5 subgroups, 4 of them not previously identified (Table S1 and Figure 6C).

**Lobula-, PVLP- and PLP-innervating uVPNs** We performed a similar analysis with uVPN groups that had dendritic innervation restricted to the lobula and axons projecting to the PVLP and posterior lateral protocerebrum (PLP) (groups IV, VI, VII and XI) (Figure S4A’). Following the same strategy, we re-clustered neurons based on the scores calculated only for the axon arbors that overlapped with the PVLP or PLP (Figure S4B). We obtained 5 distinct types, including a new subtype (Figure S4B–C and Figure S4).

**Bilateral VPNs** In addition to the analysis of the uVPNs, we also performed a hierarchical clustering of the bilateral VPNs (Figure 6A”). Of the resulting 8 distinct groups (i–viii), one matched the bilateral LC14 neurons (Otsuna and Ito, 2006).

**VPN summary** Our analysis of VPN neurons has demonstrated that similarity searches performed with neuron fragments are useful to highlight morphological features that might be important for defining neuron classes. We were able to match 11 of our defined groups to known VPN types, and furthermore described 3 new subclasses and four subtypes of uVPNs, showing that this type of analysis is able to identify new cell types even for intensively studied neuronal classes.

#### Auditory neurons

Auditory projection neurons (PNs) are characterized by their innervation of the primary or secondary auditory neuropils, the antennal mechanosensory and motor center (AMMC) and the inferior ventrolateral protocerebrum (IVLP or wedge). Several distinct types have been described based on anatomical and physiological features (Yorozu et al., 2009; Lai et al., 2012; Kamikouchi et al., 2006, 2009). Just as for the VPNs, we tested NBLAST’s ability to identify known and new cell types. In this case, we employed a two-step search strategy using a previously identified FlyCircuit neuron named by Lai et al. (2012) as the seed neuron for the first search (for details see Auditory neuron analysis). For each of the 5 types we analyzed, hierarchical clustering of the hits revealed new subtypes of known auditory PN types that differed mainly in their lateral arborizations (Figure S5E and Table S2).

#### mAL neurons

The *fruitless*-expressing mAL neurons were the first population of neurons in the brain shown to be sexually dimorphic in number and morphology (Koganezawa et al., 2010; Kimura et al., 2005). Males have around 30 neurons, females only 5. Differences in dendritic arborisations likely alter input connectivity including pheromone information detected by chemoreceptors on the male forelegs (Koganezawa et al., 2010; Kohl et al., 2015) and they regulate wing extension by males during courtship song (Koganezawa et al., 2010). We investigated whether clustering could distinguish male and female neurons and identify male subtypes. Clustering a set of 41 mAL neurons (for dataset collection see) cleanly separated male from female neurons (Figure 7A–B). Clustering analysis of the male neurons using partial skeletons that only contained the axonal and dendritic arbors (Figure 7C), identified 3 main types and 2 subtypes of male mAL neurons. The 3 types differed in the length of the ipsilateral ventral projection; this feature has previously been proposed as the basis of a qualitative classification of mAL neurons (Kimura et al., 2005). However all types and subtypes also showed reproducible differences in the exact location of their axon terminal arbors. Our analysis therefore suggests that the population of male neurons includes types with correlated differences in input-output connectivity.

**Figure 7:**
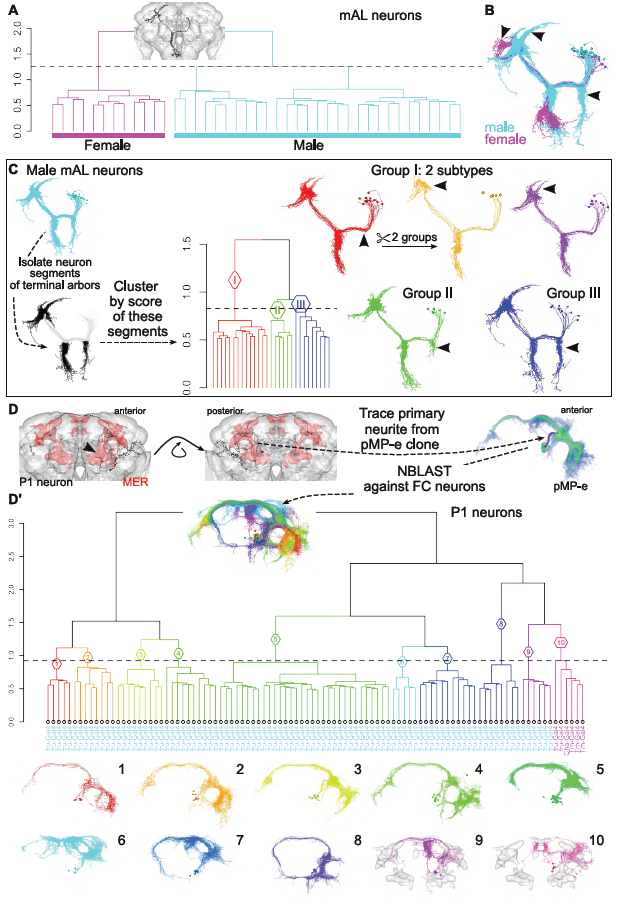
NBLAST search and classification of hits reveals subtypes of *fruitless*-expressing mAL and P1 neurons. **(A)** Analysis of the mAL neurons. Hierarchical clustering (HC) of the hits, divided into 2 groups (h=1.25). The mAL neuron used as query (fru-M-500159) is shown in the inset. The leaf labels indicate the gender of the neuron: ’F’ for female and ’M’ for male. **(B)** Neuron plot of the 2 dendrogram groups: male (in cyan) and female (in magenta). **(C)** Analysis of the male mAL neurons. The neuron segments corresponding to the terminal arbors (ipsi- and contralateral) were isolated and the neurons were clustered based on the score of these segments. HC of neurons, divided into 3 groups (groups I–III) (h=0.83). Differences in arborization indicated by arrowheads. Group I (red) can be further subdivided into two different subtypes. **(D)** Analysis of the P1 neurons. Neuron plot of a P1 neuron, fru-M-400046. The male enlarged region (MER) is shown in red. Anterior and posterior views are shown. Volume rendering of the pMP-e *fruitless* neuroblast clone, which gives rise to P1 neurons. The distinctive primary neurite was traced and used on a NBLAST search. **(D’)** HC of hits for a search against the P1 primary neurite divided into 10 groups (1–10) (h=0.92). This group of neurons corresponds to a subset of neurons obtained after a first HC analysis. Hits with a normalized score over 0.25 were collected and further selected. The inset shows a neuron plot with groups 1–10. The leaf labels show the GAL4 driver used to obtain that neuron; the colors follow the gender: cyan for male and magenta for female. Below the dendrogram, neuron plots of each group. The MER is shown in grey for groups 9 and 10.

#### P1 neurons

P1 neurons are the most significantly dimorphic *fruitless*-expressing neurons. Male P1 neurons are involved in the initiation of male courtship behavior while female P1 neurons degenerate during development due to the action of *doublesex* (Kimura et al., 2008). There are around 20 P1 neurons that develop from the pMP-e *fruitless* neuroblast clone. They have extensive bilateral arborizations in the PVLP and ring neuropil, partially overlapping with the male-specific enlarged brain regions (MER) (Kimura et al., 2008; Cachero et al., 2010) (Figure 7D).

Despite their critical role, P1 neurons have been treated as an homogeneous neuronal class. We therefore investigated whether clustering could identify anatomical subtypes. Hierarchical clustering of the P1 neuron set (for dataset collection see), distinguished 10 groups (Figure 7D’). Nine of these (1–9) contain only male *fru*-GAL4 neurons as expected. The 9 male groups have the same distinctive primary neurite and send contralateral axonal projections through the arch (Yu et al., 2010b) with extensive arborizations in the MER regions (Cachero et al., 2010). However each group shows a highly distinctive pattern of dendritic and axonal arborisations suggesting that they are likely to integrate distinct sensory inputs and to connect with distinct downstream targets.

Intriguingly, group 10 consists only of female neurons, including two neurons labeled by *fru*-GAL4. Their morphology is clearly similar to but distinct from group 9 P1 neurons, suggesting that a small population of female neurons share anatomical features (and likely originate from the same neuroblast) as these key regulators of male behavior.

### Superclusters and exemplars to organize huge data

In the previous examples we have shown that NBLAST is a powerful tool to identify known and uncover new neuron types in specific neuron superclasses. However, subsetting the dataset in order to isolate chosen neurons requires considerable time. We wished to establish a method that would organize large datasets, extracting the main types automatically, and retain information on the similarity between types and subtypes, while allowing quicker navigation. We used the affinity propagation method of clustering (Frey and Dueck, 2007), combined with hierarchical clustering to achieve this. Applying affinity propagation to the 16,129 neurons in the FlyCircuit dataset resulted in 1,052 clusters (Figure 8A–B). Using hierarchical clustering of the exemplars and by manually removing a few stray neurons, we isolated the central brain neurons (groups B–C) (Figure 8C). Another step of hierarchical clustering of central brain exemplars revealed large superclasses of neuron types (groups I–XIV), each one containing a distinct subset including, for example, central complex neurons (I), P1 neurons (II), 2 groups of KCs (γ and α′/β′ and α/β) (IV–V) and auditory neurons (VIII) (Figure 8D–D’).

**Figure 8:**
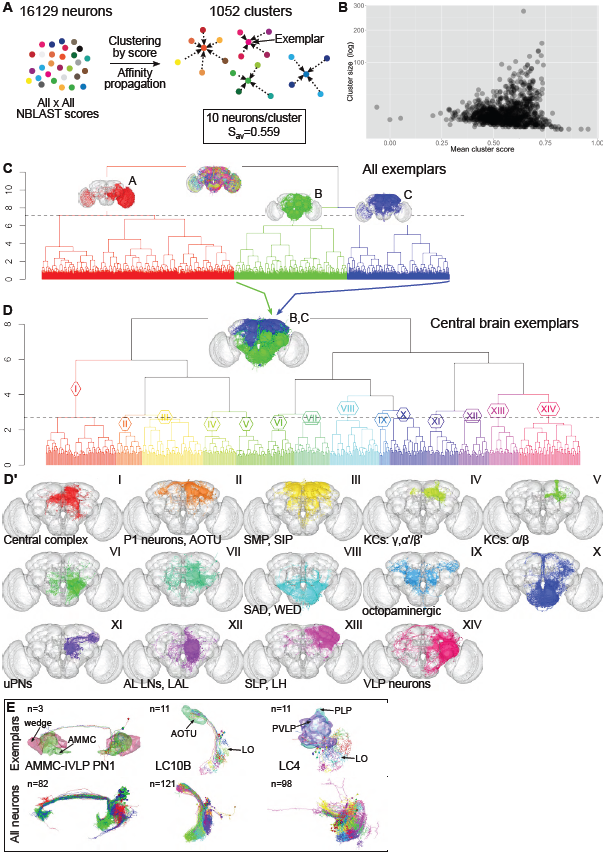
Organizing NBLAST scores by affinity propagation clustering. **(A)** Clustering by affinity propagation generated 1,052 clusters, with an average of 10 neurons per cluster and a similarity score of 0.559. Exemplars, representative neurons of each cluster, were also generated. **(B)** Plot of the mean cluster score versus cluster size. **(C)** Hierarchical clustering (HC) of the 1,052 exemplars, divided into three groups (A–C). Group A corresponds mostly to optic lobe and VPN neurons; groups B and C to central brain neurons. The insets on the dendrogram show the neurons of these groups. The main neuron types or innervated neuropils are noted. **(D)** HC of central brain exemplars (groups B and C, inset on dendrogram), divided into 14 groups, h=2.7. **(D’)** Neuron plots corresponding to the dendrogram groups in D. **(E)** Affinity propagation clusters of defined neuron types. Neuron plot of exemplars (top row) or all neurons (bottom row) for auditory AMMC-IVLP PN1 neurons (compare with Figure S5D) and VPN types LC10B (compare with Figure 6B) and LC4 (compare with Figure S4B). The number of exemplars and neurons is indicated on the top left corner. AMMC (antennal mechanosensory and motor center) in green; wedge in magenta; AOTU (anterior optic tubercle) in green; LO: lobula; PVLP (posterior ventrolateral protocerebrum) in purple; PLP (posterior lateral protocerebrum) in cyan.

The affinity propagation clusters are also useful for identifying neuronal subtypes by comparing all clusters that contain a specified neuronal type (Figure 8E). We present examples for the neuronal types AMMC-IVLP projection neuron 1 (AMMC-IVLP PN1) (Lai et al., 2012), and the uVPNs LC10B and LC4. For each of these, morphological differences are clear between clusters, suggesting that each one might help to identify distinct subtypes.

We have shown that combining affinity propagation with hierarchical clustering is an effective way to organize and explore large datasets, by condensing information into a single exemplar and by retaining the ability to move up or down in the hierarchical tree, allowing the analysis of superclasses or more detailed subtypes.

### NBLAST extensions

NBLAST is a powerful tool for working with single neurons from the adult fly, however the algorithm was designed to be general. We now illustrate NBLAST in a wide variety of experimental contexts. We first use 40 neurons reconstructed from a complete serial section electron microscopy (EM) volume of the *Drosophila* larva. Clustering NBLAST scores recovers functional groups of neurons within a multimodal escape circuit (see Figure 9A) (Ohyama et al., 2015). Pruning fine terminal branches from the EM reconstructions (mimicking light level reconstructions) has little impact on cluster assignments; therefore NBLAST clustering of coarsely skeletonized neurons could be an important step to organize electron microscopy connectome data.

**Figure 9:**
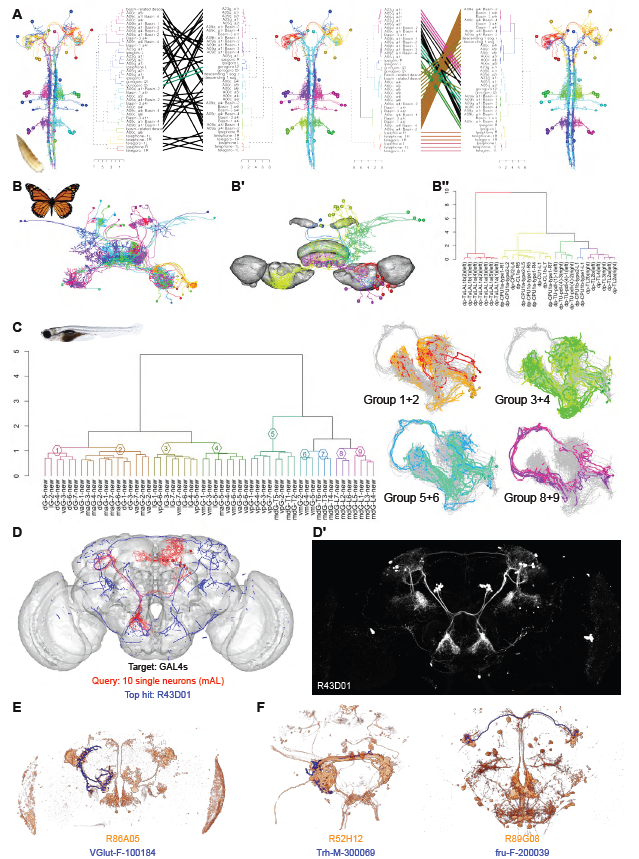
NBLAST extensions. **(A)** Larval basin goro neurons. The tanglegrams, with the dendrogram and corresponding neurons plots are shown. In the middle, the original neurons and clustering; on the left, clustering using network information and on the right, after removing the neurons first and second order terminal branches. The left tanglegram compares the network clustering to the original, whereas the right one, compares the original to the pruned clusters. **(B)** Monarch butterfly central complex neurons. Plot of all neurons. **(B’)** Neurons after mirroring and clustering. The colors correspond to the dendrogram groups. A few of of the brain neuropils are shown in grey. **(B”)** Hierarchical clustering after mirroring neurons. **(C)** Zebrafish mitral neurons. On the left, hierarchical clustering after mirroring neurons. On the right, 4 examples of pairwaise comparisons of neuron groups. The colors correspond to the dendrogram groups. **(D)** Dotprops representation of R43D01 expression pattern (blue) with best matched FlyCircuit mAL neurons (red). **(D’)** R43D01 expression pattern. **(E)** Candidate V-glomerulus-selective GAL4 line expression pattern with single neuron (VGlut-F-100184). **(F)** Identified GAL4 line expression patterns found with AMMC-AMMC PN1 auditory neuron (Trh-M-300069) and an AOTUv2 lineage neuron (fru-F-200039).

We next show two examples applying NBLAST to single cell data from another invertebrate, the monarch butterfly, and a vertebrate, the larval zebrafish (Figure 9B, C and). Clustering 29 monarch butterfly neurons from the central complex (Heinze et al., 2013) largely matches neuronal types defined by expert neuroanatomists – the few discrepancies were reviewed with the data provider (S. Heinze, personal communication) and determined to be cases where computationally defined cell groups revealed features that were orthogonal to expert classification but still a valid classification.

The zebrafish data consisted of 55 mitral cells (second order olfactory neurons) projecting to a variety of higher brain areas (Miyasaka et al., 2014). NBLAST clustering identified clearly distinct morphological groups (Figure 9C). Very similar neurons were co-clustered both by our algorithm and that of the original authors, but clustering of distantly related neurons was distinct. Only future experiments will show if one clustering has more functional relevance.

In our final example,we apply NBLAST to a distinct but experimentally vital form of neuroanatomical image data. Circuit neuroscience in many model organisms depends on manipulating circuit components with cell type specific driver lines. We have registered (Manton et al., 2014) and processed image data from the most widely used *Drosophila* collection, 3501 GMR driver lines generated at the Janelia Research Campus (Jenett et al., 2012). We applied an image processing pipeline emphasizing tubular features (Masse et al., 2012), generating a vector cloud representation identical to that used elsewhere in this paper. These data (9Gb for 3501 image stacks) can be queried with single neurons or tracings in less than 30 seconds on a desktop computer. To demonstrate this approach at scale we mapped these GAL4 data to the same template space (Manton et al., 2014) as the FlyCircuit single neurons and computed NBLAST scores for 16129 neurons against 3501 driver lines. We provide a simple web server for these queries at jefferislab.org/si/nblast/on-the-fly. We showcase this by identifying GAL4 driver lines targeting the sexually dimorphic mAL neuron population (Figure 9D, D’, see also 7). We selected 10 mAL neurons and then examined the 10 GAL4 lines with the highest mean scores. The top hit line R43D01 has just been identified as targeting this population (Kallman et al., 2015) and all the top 10 hits target the same population.

As a second example, we looked at driver lines labeling olfactory projection neurons targeting the CO_2_-responsive V glomerulus. Comprehensive single cell labeling identified classes critical for behavioral responses to different CO_2_concentration (Lin et al., 2013). However one class highly selective for the V glomerulus could not be functionally studied because no GAL4 line was identified. Searching this neuron we found the fourth hit (R86A05) was highly selective for this cell type (Figure 9E). Finally, we take an auditory interneuron (AMMC-AMMC PN1, see S5 and S2) and a presumptive visual interneuron of the anterior optic tubercle. The top 10 hits for both neurons included numerous matching GAL4 lines; we display one example for each in Figure 9F. Although all of these lines label multiple neuronal classes, NBLAST enables very rapid identification of lines containing a neuronal population of interest that could be used for the construction of completely cell-type specific lines by intersectional approaches (Luan et al., 2006).

## Discussion

The challenge of comprehensive mapping neuronal types in the brain depends on establishing methods that facilitate unbiased identification of neuron types from pools of thousands or millions of individual neurons. Comparison of neurons relies on morphology and brain position, essential determinants of their function and synaptic partners. A neuron search algorithm should be: (1) accurate, generating biologically meaningful hits; (2) computationally inexpensive; (3) enable interactive searches for data exploration (4) generally applicable. We have described NBLAST, a neuron search algorithm that satisfies all these criteria.

First, the algorithm correctly distinguishes closely related subtypes across a range of major neuron groups, achieving 97.6 % accuracy for 35 classes of olfactory projection neuron. Unsupervised neuron clustering based on NBLAST scores, correctly organized neurons into known types. We did find that neuron sizes (especially when very small) can influence the algorithm: one future research area will be to convert the raw scores that we have used into an expectation (E) value (cf. BLAST), that would directly account for the size of a neuron and the database.

Second, NBLAST searches are very fast, with pairwise comparisons taking about 2 ms on a laptop. Furthermore, for defined datasets all-by-all scores can be pre-computed allowing immediate retrieval of NBLAST scores for highly interactive analysis. With data volumes increasing, strategies to query and store these data need to be investigated. One effective approach to handle much larger numbers of neurons will be to compute sparse similarity matrices, storing the top *n* hits for a given neuron. Alternatively, queries could be computed only against the non-redundant set of neurons that collectively embody the structure of the brain (similar to the strategy employed by UniProt (Suzek et al., 2007)). For the fly brain this could not exceed 50,000 neurons (due to strong bilateral symmetry) and we expect the actual number to be < 5,000. Our clustering of all ~16,000 neurons of the FlyCircuit dataset identified ~1,000 exemplars providing a non-redundant data set that we use for rapid searches.

Third NBLAST permits multiple search strategies and types of analysis. Searches can use neuron fragments, or tracings from complex image data as queries and databases of GAL4 lines as targets. Closely related neuronal types can be distinguished by clustering of only their terminal arbors without considering common features such as axon tracts.

Finally, one important question is the extent to which our approach generalizes. This issue largely reduces to the relationship between length scales of neurons being examined and their absolute spatial stereotypy. Our method implicitly assumes spatial co-localization of related neurons; this is enforced by the use of image registration. Our strategy should be appropriate for any situation in which neuronal organization is highly stereotyped at the length scale of the neurons themselves. There is already strong evidence that this is true across large parts of the brain for simple vertebrate models like the larval zebrafish; indeed we show that our NBLAST method can be applied directly to olfactory projectome data (Miyasaka et al., 2014). Mouse gene expression (Lein et al., 2007) and long range connectivity also show global spatial stereotypy as evidenced by recent atlas studies combining sparse labeling and image registration (Zingg et al., 2014; Susaki et al., 2014; Oh et al., 2014)). Our method could be adapted for querying and hierarchical organization of these datasets by calculating an appropriate scoring matrix.

However there are situations in which global brain registration is not appropriate For example the vertebrate retina has both laminar and tangential organization. Sümbül et al. (2014) recently introduced a registration strategy that showed that lamination of retinal ganglion cells is spatially stereotyped to the nearest micron. However retinal interneurons and ganglion cells are organized in mosaics tangential to the retinal surface; global registration is not appropriate in this plane. The situation is similar for parallel columns of the outer *Drosophila* optic lobe. We see two possible approaches: local re-registration, mapping neurons onto a single canonical column or, amassing sufficient data so that neurons from neighboring columns/mosaics tile the brain, enabling identification of related groups by clustering or graph theoretic approaches.

Cataloguing all neuron types in the brain will rely not only on effective measures of neuronal similarity, but also on simple, unbiased methods for automated classification of neurons into functionally relevant types. This is a challenging problem: it may be necessary to combine morphological approaches with other data such as connectivity patterns, single neuron gene expression patterns or physiological properties to provide unambiguous automated classification (reviwed by Armañanzas and Ascoli, 2015). We have shown that NBLAST scores define a highly effective similarity metric that can be combined with hierarchical clustering and a specific dendrogram cut height to define a very wide range of neuronal classes. This approach enables very rapid exploratory analysis of new cell types even without expert neuroanatomical knowledge. Indeed for *Drosophila* neurons it seems that NBLAST clustering is sufficient to define cell type.

Sümbül et al. (2014) recently explored the issue of defining the optimal dendrogram cut height for morphological clustering of 363 mouse retinal ganglion cells, establishing a reliable approach for these specific neurons. Nevertheless, our experience from the 16,129 neuron validation set is that differences in similarity levels within classically defined neuronal types preclude the existence of a universal value for dendrogram cut height. Some of this range (0.7 to 2 in this study) is probably due to differences in definitions: classic neuronal types may in some cases require splitting for consistency – we see evidence for this in the Kenyon cell and visual projection neuron datasets. More sophisticated statistical criteria may enable automated classification, especially when combined with measurements of e.g. physiological or gene expression data (Armañanzas and Ascoli, 2015). However all approaches to defining cluster numbers (i.e. statistically-based cell types) depend on biological priors that must be acknowledged. Nevertheless NBLAST’s speed and sensitivity and the size of this validation dataset represent a significant step towards fully automated classification.

Finally we note that NBLAST can identify genetic driver lines labeling a given query neuron. The pre-computed NBLAST result matrix that we provide for the GMR GAL4 collection will be of immediate utility to our *Drosophila* colleagues planning experimental studies of particular cell classes. Thus NBLAST can provide a vital link between studies of anatomical logic and neural circuit function.

## Experimental Procedures

### Image Preprocessing

flycircuit.twsupplied 16,226 raw confocal stacks in Zeiss LSM format on a single 2 TB hard drive in April 2011. Each stack was uncompressed, then read into Fiji/ImageJ (http://fiji.sc/Fiji) where channels were split and resaved as individual NRRD (http://teem.sourceforge.net/nrrd) files. Where calibration information was missing in the LSM metadata, we used a voxel size of (0.318427, 0.318427, 1.00935) microns as recommended by the FlyCircuit team. There were two issues to solve before images could be used for registration: 1) identifying which image channel contained the anti-Dlg (discs large 1) counterstaining 2) determining whether the brains had been imaged from anterior to posterior, or the reverse. The first issue was solved by exporting metadata associated with each LSM using the LOCI bioformats (http://loci.wisc.edu/software/bio-formats) plugin and developing heuristics to automate the identification of the channel sequence; for a minority of images metadata was missing and channel order was determined manually. Slice order could not be determined automatically from the image metadata. We therefore made maximum intensity Z projections (using the unu tool, http://teem.sourceforge.net/unrrdu) for the labeled neuron channel for each stack. Each projection was then compared with the matching thumbnail available from the flycircuit.tw website. The correlation score between the projection and thumbnail images was calculated both with and without a mirror flip across the YZ plane; a large correlation score for only one orientation was used as evidence for a given slice ordering. A small number of ambiguous results were verified manually. We successfully preprocessed 16,204/16,226 total images i.e. a 0.14 % failure rate. 12 failures were due to mismatches that could not be resolved between the segmented neuron present in the LSM file and the thumbnail image on the flycircuit.tw website; the remaining 10 failures were due to physical offsets between the brain and GFP channels or corrupt image data.

### Template Brain

The template brain (FCWB) was constructed by screening for whole brains within the FlyCircuit dataset and manually selecting a pool of stacks that appeared of good quality. Separate average female and average male template brains were constructed from 17 and 9 brains, respectively using the CMTK (http://www.nitrc.org/projects/cmtk) avg_adm tool which takes a single brain as a seed. After five iterations the resultant average male and average female brains were placed in an affine symmetric position within their image stacks so that a simple horizontal (*x*-axis) flip of either template brain resulted in an almost perfect overlap of left and right hemispheres. Finally the two sex-specific template brains were then averaged (with equal weight) to make an intersex template brain using avg_adm. Since this template was designed to provide an optimized registration target for the FlyCircuit dataset, no correction was made for the disparity between the XY and Z voxel dimensions common to all images in the dataset. The scripts used for the construction of the template are available at https://github.com/jefferislab/MakeAverageBrain.

### Image Registration

Image registration of Dlg neuropil staining used a fully automatic intensity-based (landmark free) 3D image registration implemented in the CMTK toolkit (Rohlfing and Maurer, 2003; Jefferis et al., 2007). An initial linear registration with 9 degrees of freedom (translation, rotation and scaling of each axis) was followed by a non-rigid registration that allows different brain regions to move independently, subject to a smoothness penalty (Rueckert et al., 1999). It is our experience that obtaining a satisfactory initial linear registration is crucial. All registrations were therefore manually checked by comparing the reformatted brain with the template in Amira (academic version, Zuse Institute, Berlin), using ResultViewer https://bitbucket.org/jefferis/resultviewer. This identified about 10 % of brains with poor initial registrations. For these images a new affine registration was calculated using a Global Hough Transform (Ballard, 1981; Khoshelham, 2007) with an Amira extension module available from 1000shapes GmbH; the result of this affine transform was again manually inspected. In the minority of cases where this approach failed, a surface based alignment was calculated in Amira after manually aligning the two brains. Following satisfactory initial affine registration, a non-rigid registration was calculated. Finally each registration was checked manually in Amira against the template brain. This sequential procedure was resulted in successfully registration of 16,129/16,204 preprocessed images, giving a registration failure rate of 0.46 %.

### Image Postprocessing

The confocal stack for each neuron available at http://flycircuit.tw includes an 8 bit image containing a single (semiautomatically) segmented neuron prepared by Chiang et al. (2011). This image was downsampled by a factor of 2 in *x* and *y*, binarized with a threshold of 1 and skeletonized using the Fiji plugin ‘Skeletonize (2D/3D)’ (Doube et al., 2010). Dot properties for each skeleton Masse et al. (2012), were extracted using the dotprops function of our new nat package for R. This converted each skeleton into segments, described by its location and tangent vector. Neurons on the right side of the brain were flipped to the left by applying a mirroring and a flipping registration as described in Manton et al. (2014). The decision to flip a neuron depended on earlier assignment of each neuron to a brain hemisphere using a combination of automated and manual approaches. Neurons whose cell bodies were more than 20 μm away from the mid-sagittal YZ plane were automatically defined as belonging to the left or right hemisphere. Neurons with cell bodies inside this 40 μm central corridor were manually assigned to a hemisphere, based on cell body position (right or left side), path taken by the primary neurite, location and length of first branching neurite. For example, neurons that had a cell body on the midline with significant innervation from the first branching neurite near the cell body on the left hemisphere, with the rest of the arborization on the right, were assigned to the left side and not flipped. The cell body positions used were based on those published on the http://flycircuit.tw website for each neuron; these positions are in the space of the FlyCircuit female and male template brains (typical_brain_female and typical_brain_male). In order to transform them into the new FCWB template that we constructed, affine bridging registrations were constructed from the typical_brain_female and typical_brain_male brains to FCWB and the cell body positions were then transformed to this new space. Since these cell body positions depend on two affine registrations (one conducted by Chiang et al. (2011) to register each sample brain onto either their typical_brain_female or typical_brain_male templates and a second carried out by us to map those template brains onto our FCWB template) these positions are likely accurate only to ±5 microns in each axis.

### Neuron Search

The neuron search algorithm is described in detail in Results and Figure 1. The reference implementation that we have written is the nblast function in the R package nat.nblast, which depends on our nat package (Jefferis and Manton, 2014). Fast nearest neighbor search, an essential primitive for the algorithm uses the nabor package (Jefferis, 2015), a wrapper for the nabo C++ library (Elseberg et al., 2012). The scoring matrix that we used for FlyCircuit neurons was constructed by taking 150 DL2 projection neurons, defining a neuron type at the finest level, and calculating the joint histogram of distance and absolute dot product for the 150 × 149 combinations of neurons, resulting in 1.4 × 10^7^ measurement pairs; the number of counts in the histogram was then normalized (dividing by 1.4 × 10^7^) to give a probability density, *p*_match_. We then carried out a similar procedure for 5,000 random pairs of neurons sampled from the FlyCircuit dataset to give *p*_rand_. Finally the scoring matrix was calculated as 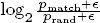 where ε (a pseudocount to avoid infinite values) was set to 1 × 10^-6^.

### Clustering

We employed two different methods for clustering based on normalized NBLAST scores. We used Ward’s method for hierarchical clustering, using the default implementation in the R function hclust. This method minimizes the total within-cluster variance, and at each step the pair of entities or clusters with the minimum distance between clusters are merged (Ward Jr, 1963). The resulting dendrograms were cut at a single selected height chosen for each case. This value is shown as a dashed line in all dendrograms. By default, R plots the square of the Euclidean distance as the *y* axis, but in the plots shown, the height of the dendrogram corresponds to the unsquared distance.

For the analysis of the whole dataset, we used the affinity propagation method. This is an iterative method which finds exemplars which are representative members of each cluster and does not require any *a priori* input on the final number of clusters (Frey and Dueck, 2007) as implemented in the R package apcluster (Bodenhofer et al., 2011). The input preference parameter (*p*) can be set before running the clustering. This parameter reflects the tendency of data samples to become an exemplar, and controls the final number of clusters. We used *p* = 0, since this is the value where on average matched segments are equally likely to have come from matching and non-matching neurons. Empirically this parameter produced clusters that mostly grouped neurons of the same type according to biological expert opinion.

### Neuron Tracing

Neuron tracing was carried out in Amira (commercial version, FEI Visualization Sciences Group, Merignac, France) using the hxskeletonize plugin (Evers et al., 2005) or in Vaa3D (Peng et al., 2014) on registered image data. Traces were loaded into R using the nat package. When necessary, they were transformed into the space of the FCWB template brain Manton et al. (2014).

### Computer Code and Data

The image processing pipeline used two custom packages for the R statistical environment (http://www.r-project.org): https://github.com/jefferis/nat and https://github.com/jefferis/nat.as that coordinated processing by the low level registration (CMTK) and image processing (Fiji, unu) software mentioned above. NBLAST neuron search is implemented in R package nat.nblast available at https://github.com/jefferislab/nat.nblast. Analysis code specific to the fly-circuit dataset is available in a dedicated R package https://github.com/jefferis/flycircuit, with a package vignette showcasing the main tools that we have developed. Further details of these supplemental software and the associated data are presented at http://jefferislab.org/si/nblast. The registered image dataset can be viewed in the stack viewer of the http://virtualflybrain.org website and all 16,129 registered single neuron images and corresponding skeletonized neurons will be available at http://jefferislab.org/si/nblast or on request to GSXEJ on a hard drive; the unregistered data remain available at http://flycircuit.tw.

Precomputed score matrices of all by all FlyCircuit neurons and all FlyCircuit neurons against all GMR GAL4 lines are distributed through the https://github.com/jefferis/flynblastscores R package or available via a remote web service that can return the top n hits for any neuron (see vfb_nblast function in package https://github.com/jefferis/vfbr). Finally a fully interactive web application is available by following the link http://bit.ly/nblast.

## Acknowledgments

We first of all acknowledge the flycircuit.tw team for generously providing the raw image data associated with Chiang et al. (2011). Images from FlyCircuit were obtained from the NCHC (National Center for High-performance Computing) and NTHU (National Tsing Hua University), Hsinchu, Taiwan. We thank Stanley Heinze for sharing monarch butterfly neuron tracings and discussions about cell classification. We thank Albert Cardona and Marta Zlatic for providing larval *Drosophila* electron microscopy skeleton data. We thank Nobuhiko Miyasaka, Yoshihiro Yoshihara, Ignacio Arganda-Carreras, Uygar Sümbül and Sebastian Seung for sharing zebrafish mitral cell reconstructions. We thank members of the Jefferis lab for comments on the manuscript, Jake Grimmett and Toby Darling for their ongoing assistance with the LMB’s compute cluster and Torsten Rohlfing for discussions about image analysis and registration. We thank the Virtual Fly Brain project for their help in linking and incorporating some of the results of this study in the http://virtualflybrain.org website.

This study made use of the Computational Morphometry Toolkit, supported by the National Institute of Biomedical Imaging and Bioengineering. This work was supported by the Medical Research Council [MRC file reference U105188491] and European Research Council Starting Investigator and Consolidator Grants to GSXEJ, who is an EMBO Young Investigator.

## Supplemental Information

### Supplemental Experimental Procedures and Results

#### Algorithm Design Process

The algorithm design process was primarily motivated by a requirement for rapid and sensitive searches of neuron databases. It was necessary to consider both the data structure and the similarity algorithm jointly in the face of the design requirements.

Our first application would be data acquired in *Drosophila*, where previous studies using image registration have shown a high degree of spatial stereotypy (standard deviation of landmarks ~2.5 μm in each axis for a brain of 600 μm in its longest axis, Jefferis et al., 2007). Therefore one key design decision was to use co-registered data rather than calculating similarity using features of the neuron that are independent of absolute spatial location.

On the algorithm side, the key initial design decision was whether to develop a direct pairwise comparison algorithm or to use a form of dimensional reduction to map neuronal structure into a lower-dimensional space. The major advantage of the latter approach is that the similarity between neurons can be computed directly and almost instantaneously in the low dimensional space. However, the construction of a suitable embedding function either requires existing knowledge of neuronal similarity (likely supplied by experts in the form of large amounts of training data), huge amounts of unlabeled data that enable direct learning of features (e.g. Le et al., 2012), or a strategy based on successful extraction of key image features.

A number of considerations made us favor the approach of direct pairwise comparison. First, we suspected that it would be possible to make a more sensitive algorithm by working with the original data. Second, the amounts of image data available did not seem large enough to avoid a requirement for extensive labeled training data. Third, we reasoned that our own intuitions about neuronal similarity could be better expressed in the original physical space of the neuron than in a low dimensional embedding. Our own exploratory analysis in which we summarized each neuron in different ways as feature vectors of the same dimension and used a comparison function in the feature space (SP, GSXEJ unpublished observations) confirmed that constructing a sensitive metric of this sort is challenging.

The selection of a pairwise similarity metric meant that we had to give particularly careful consideration to performance issues in the design phase. We set two practical performance targets: 1) being able to carry out searches of a single neuron against a database of 10,000 neurons in less than a minute on a simple desktop or laptop computer. 2) Being able to complete all-against-all searches for 10,000 neurons (10^8^ comparisons) in < 1 day on a powerful desktop computer. These targets meant that each elementary comparison operation should take around 5 ms or less. Image pre-processing carried out once per neuron would therefore be a good investment if it reduced the time taken for each pairwise comparison. These considerations prompted us to generate a spatially registered, compact representation of each neuron as a separate pre-processing step, rather than develop an algorithm that simultaneously solved both the spatial alignment and similarity problem.

#### Algorithm Scoring

As described in the main results section, we defined NBLAST raw scores as the sum of segment pair scores:

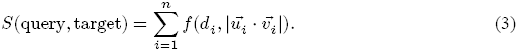

We initially experimented with a function based on expert intuition:

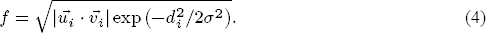

This includes a negative exponential function of distance (related to the normal distribution), with a free parameter *σ* based on our previous estimates of the variability in position within the fly brain of landmarks after registration (Jefferis et al., 2007; Yu et al., 2010b) set to 3 μm. Although this provided a useful starting point, we were unhappy with a scoring system that required parameters to be specified rather than derived empirically from data. This then motivated us to investigate the statistical scoring approach described in based on log probability ratios. This single parameter approach may still be helpful in some situations where insufficient neurons are available in order to define a statistical scoring matrix (we used 150 similar neurons and 5,000 random pairs). It can also enable a bootstrapping approach for new datasets in order to help identify similar pairs of neurons that can then be used to define a full scoring matrix.

#### Kenyon cell analysis

**Dataset collection and dividing into types** We collected the dataset of initial KCs by performing a forward and reverse search against all neurons using one identified KC (fru-M-500225). We then selected neurons that had both raw scores above -2,500 (2,088 neurons). We performed affinity propagation clustering (Frey and Dueck, 2007) of these neurons, obtaining 59 clusters, and manually verified each one, resulting in 1,562 neurons being identified as KCs. An additional search for high scorers against these KC exemplars uncovered an extra 102 neurons, bringing the total number of KCs used in our analysis to 1,664, representing 10.3 % of the FlyCircuit dataset.

We performed hierarchical clustering of the KCs, based on the NBLAST scores, and divided the dendrogram into two groups (Figure 4A). Contrary to expectations, one group contained both the γ and α′/β′ neurons, whereas the other group consisted exclusively of α/β neurons (Figure 4B–D), the largest subset in our sample. We separated α′/β′ from the γ neurons in a subsequent hierarchical clustering of this group. We performed additional analysis for each of the neuron types.

**Analysis of γ neurons** Hierarchical clustering of the 470 γ neurons resulted in a dendrogram which we divided into three groups (I–III) (Figure 4B’). The number of clusters was chosen by visual inspection in order to reveal differences in morphology and organization between the groups. Groups I and III corresponded to the classical γ neurons while group II matched atypical γ neurons. There were differences in neurite positioning in the calyx, from medial to lateral, with group I being the most medial, followed by groups II and III. There were also differences in the gamma lobe, with group II occupying the anterodorsal region, while groups I and III were mostly mixed in the rest of the lobe. A subsequent clustering analysis of the classical γ neurons divided into 4 groups (groups A-D) revealed that there were differences between the groups in their medial to lateral position in the calyx. These differences correlated to a certain degree with differences in the dorsal/ventral position of the projections in the γ lobe, with groups C and D, the most medial, being also the most dorsal (Figure 4B”). These observations suggest that the relative position of the projections of classical γ neurons is maintained at the calyx and γ lobe.

In order to understand if the relative position of the classical γ neurites was maintained in between the calyx and the γ lobe, we clustered the neurons based on the scores of the segments in the peduncle. We took neuron skeletons from classical γ neurons and isolated the axon arbors that co-localized with the peduncle volume (Figure S3B’). We then carried out a new clustering based on all-by-all NBLAST scores of these partial skeletons, cutting the dendrogram at a level defined by visual inspection (4 groups). The overall organization almost fully recapitulated the positioning of the neurites in the whole neuron analysis (compare Figure 4B’–B” with Figure S3B’). A clear and expected lamination was found in the peduncle, with neurites occupying the most outer stratum. Differences in the medial to lateral positioning of neurites in the calyx followed the previously observed organization, with the most medial groups occupying the dorsal region of the gamma lobe. The overall organization almost fully recapitulated the positioning of the neurites in the whole neuron analysis (for more information see Figure S3 and Supplemental Results). Thus, the stereotypical organization of the classical γ neurons is maintained throughout the neuropil.

Group II of the γ neurons matched atypical γ neurons (γ dorsal neurons) (Aso et al., 2014) with neurons that extended neurites posteriolaterally in the calyx and projected to the most dorsal region of the γ lobe (Figure 4B”’). Hierarchical clustering of these neurons resulted in a dendrogram that we divided into 3 groups (a-c). This number of groups isolated the previously identified subtype of atypical γ neurons – γd neurons (Aso et al., 2009) – into one group (group a). These neurons extend neurites ventrolaterally at the level of the calyx (identified as ventral accessory calyx) (Aso et al., 2014). The other 2 groups (b, c) correspond to uncharacterized types. Although they project to a similar region in the γ lobe, their dendrites do not extend laterally and their calyx neurites are longer than γd neurons.

**Analysis of α′/β′ neurons** Hierarchical clustering analysis of the α′/β′ neurons and separation into 4 groups highlighted the characterized subtypes of α′/β′ neurons (Figure 4C–C’). They differ in their anterior/posterior position in the peduncle and β′ lobe with three types described – α′/β′ anterior and posterior (α′/β′ap) and α′/β′ medial (α′/β′m) (Tanaka et al., 2008, Y. Aso, personal communication)). Although we were unable to unambiguously assign a α′/β′ subtype to each group i–iv because our sample was too small, there were clear trends. Neurites of neurons in groups i and iv were more anterior than the other 2 groups (ii, iii) in both the peduncle and β′ lobe. These relative positions were not maintained in the calyx, with the two the anterior groups (i, iv) occupying either a medial or a lateral position.

**Analysis of α/β neurons** The largest subset of KCs corresponds to α/β neurons (Figure 4D). During the analysis of this group we found 18 neurons that did not correspond to α/β cells, since they innervated either only the β or the α lobes, and they were removed from the analysis. We performed hierarchical clustering on the remaining 1,091 cells and divided the resulting dendrogram into four groups (1–4) (Figure 4D’), which matched the four neuroblast lineages from which they originate (Zhu et al., 2003). The relative position of the neurites of the four groups within the calyx is somewhat maintained in the peduncle, with the lateral neuroblast clones (group 1 AL; group 2 PL) extending along the dorsolateral peduncle, while the medial clones (group 3 AM; group 4 PM) occupy a more ventromedial region. Hierarchical clustering of each group revealed an expected common organization for all neuroblast clones (Figure 4D” and Figure S3A). For all groups there was a clear distinction between the late born core (α/β core, α/β-c) and early born peripheral neurons (α/β surface, α/β-s). Core neurons are on the inside stratum of the α lobe. They are also reported to occupy the inside stratum of the peduncle and β lobe (Tanaka et al., 2008). We were unable to observe this, although the projections of α/β core neurons were ventral to α/β surface neurons in both the peduncle and β lobe. There was also a trend for α/β surface neurons to occupy a more medial position in the calyx in comparison to α/β core ones. A subgroup of group 2 corresponded to the α/β posterior or pioneers neurons (α/βp). The α/βp neurons are the earliest born α/β and they innervate the accessory calyx, run along the surface of the posterior peduncle into the β lobe but stop before reaching the medial tip (Tanaka et al., 2008). A new clustering based on peduncle position of the neuron segments did not recapitulate the relative positions of the calyx neurites for each of the neuroblast clones observed in the whole neuron analysis suggesting that the relative position of the α/β neurons in the peduncle does not completely reflect their stereotypical organization in the calyx (for more information see Figure S3 and Supplemental Results).

In order to investigate the stereotypical organization of α/β neurites, we performed a similar analysis as for the classic γ neurons, isolating the axon arbors that co-localized with the peduncle for groups 1 to 4 (Figure S3B”). The new clustering based on peduncle position of these partial neuron skeletons did not recapitulate the relative positions of the calyx neurites for each of the neuroblast clones observed in the whole neuron analysis (compare Figure 4D’ with Figure S3B”). In addition, there was no clear organization of neurites in the α lobe that correlated with their position in the peduncle.

#### Olfactory projection neuron analysis

We started by manually classifying the 400 uPNs in the FlyCircuit dataset by glomerulus, neuroblast lineage, and axon tract, using the original image stacks. The definition of the manual gold standard annotations was an iterative process that took several days. The first round accuracy was about 95 %. Numerous discrepancies were revealed by subsequent NBLAST analysis and difficult cases were resolved by discussion between two expert annotators before finalizing assignments. We excluded 3 neurons for which no conclusion could be reached. We found a very large number of DL2 uPNs, 145 DL2d and 37 DL2v neurons, in a total of 397 neurons. Nevertheless, our final set of uPNs broadly represents the total variability of described classes and contains neurons innervating 35 out of 56 different glomeruli (Tanaka et al., 2012), examples of the three main lineage clones (adPNs, lPNs and vPNs) in addition to one bilateral uPN, and neurons that follow each of the three main tracts (medial, mediolateral and lateral antennal lobe tracts). For subsequent analysis, we removed 3 neurons for which registration failed.

#### Visual projection neuron analysis

We started with the 1,052 exemplars found by affinity propagation clustering of NBLAST scores (Figure 8). We then clustered those exemplars using hierarchical clustering and found that extrinsic and intrinsic optic lobe neurons together formed a distinct “optic lobe” group within this (Figure 8C). We then collected all neurons associated with those “optic lobe” exemplars and calculated the overlap of neurons with each of the standard neuropils defined by Ito et al. (2014) (see Experimental Procedures and Manton et al., 2014 for technical details). This then allowed us to separate neurons by innervation pattern into 3 groups: 1) ipsilateral optic lobe neuropils only (see Figure S4D), 2) ipsilateral and central brain neuropils (unilateral VPNs, uVPNs) or 3) both optic lobes and central brain neuropils (bilateral VPNs). This selection procedure resulted in a set of 1,793 uVPNs and 72 bilateral VPNs.

#### Auditory neuron analysis

We employed a two-step search strategy. First, for 5 of the auditory types, we used the FlyCircuit neuron named by Lai et al. (2012) as the seed neuron for the first search. Candidate neurons were selected using strict anatomical criteria. A second search was then done using these candidates as query neurons and collecting all high scorers (score over 0.5). These neurons corresponded to our set for each of the types.

#### mAL neuron analysis

The set of maL neurons resulted from a search with a seed mAL neuron, fru-M-500159. We then collected 41 hits with a mean NBLAST score greater than 0.2.

#### P1 neuron analysis

The set of P1 neurons was identified by searching the FlyCircuit dataset with a tracing of the distinctive primary neurite of a pMP-e clone (Cachero et al., 2010). Hierarchical clustering of the top hits, after manual verification, identified a subset consisting solely of P1 neurons which was used in the subsequent analysis.

#### A.1.9 NBLAST extension analysis

The 29 monarch butterfly neurons were scaled by a factor of 5 before clustering. The 55 zebrafish mitral cells where mapped onto the right hand side of the larval fish brain before clustering.

### Online resources

The online resources provided in the paper are listed at http://jefferislab.org/si/nblast. In addition to the open source software described in the Experimental Procedures we also provide:

- Code and instructions to generate some of the figure panels used in the paper. Instructions can be found at https://github.com/jefferislab/NBLAST_figures. A video demo is also available at http://youtu.be/LJgZejabqqg.
- The affinity propagation clustering of the flycircuit.tw dataset (as in Figure 8D), excluding intrinsic optic lobe neurons. This can be viewed online at http://jefferislab.org/si/nblast/clusters, including interactive 3D rendering of clusters powered by WebGL.
- The clusters identified by affinity propagation clustering (including intrinsic optic lobe neurons) are all indexed by the Virtual Fly Brain (http://www.virtualflybrain.org, VFB) website, which links to 3D WebGL renderings of each cluster hosted at jefferislab.org. Clusters can be identified by VFB queries for the neuropil region that they innervate. For example, search from the VFB homepage for “AMMC”. From the [results page](http://www.virtualflybrain.org/site/tools/anatomy_finder/index.htm?id=FBbt:00003982&name=AMMC%20(synonym)), choose the query “Images of neurons with: some part here (clustered by shape)”. A list of clusters, with thumbnail images is [displayed](http://www.virtualflybrain.org/do/cluster_list.html?action=cluster_found&id=FBbt:00003982); single exemplars are also displayed for each cluster, hyperlinked to the original data at flycircuit.tw. The images of the individual neurons that are part of this cluster can also be displayed in the stack browser from this page (“List individual members” or “Open all in viewer”). Clicking on a cluster thumbnail links to a page which includes a [snapshot](http://flybrain.mrc-lmb.cam.ac.uk/vfb/fc/clusterv/3/fru-F-800038/) and [3D rendering](http://flybrain.mrc-lmb.cam.ac.uk/vfb/fc/clusterv/3/fru-F-800038/webgl.shtml) of the cluster, and information about the neurons that are part of this cluster, including links to the appropriate Neuron ID pages at flycircuit.tw; a second table provides links to result pages for the most similar clusters. A video demo is available at http://youtu.be/YFsxjkdr5y8.
- An online web-app allowing on-the-fly NBLAST queries of FlyCircuit neurons against other FlyCircuit neurons, as well as queries of user-uploaded neurons against the FlyCircuit dataset, available at http://jefferislab.org/si/nblast/on-the-fly/.
- Video demos for the online resources are available at http://jefferislab.org/si/nblast/www/demos.

### Supplemental Data

**Figure S1:**
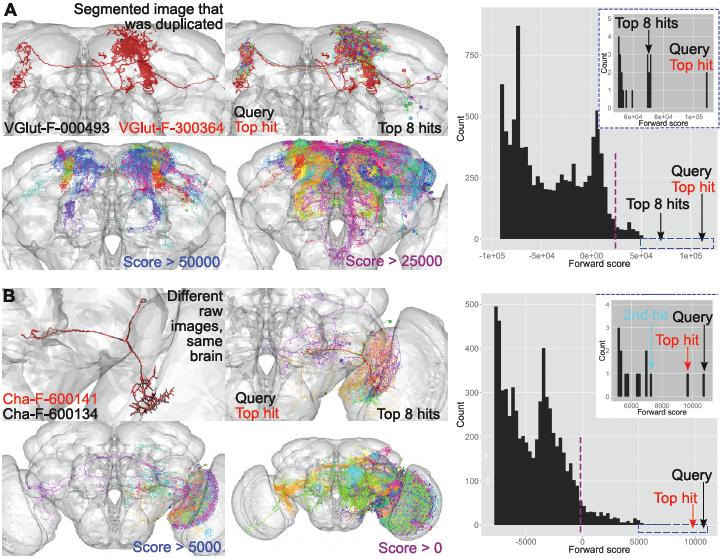
Neuron search with NBLAST, related to Figure 3. **(A)** NBLAST search with VGlut-F-000493 as query. Neuron plots of (from higher to lower score): the query (black) and top hit (red), top 8 hits, hits with a score over 50,000 and hits with a score over 25,000. The top hit corresponds to a segmented image that was duplicated. It perfectly overlays the query neuron. As the score decreases, so does the similarity of the hits to the query. On the right, histogram of forward scores. Only hits with scores over -100,000 are shown. The score of the query, top hit and top 8 hits are indicated. A dashed purple line marks 25,000. The left inset shows a zoomed view of the top hits (score > 50,000) (dashed blue rectangle in main plot). The score of the query, top hit and top 8 hits are indicated. **(B)** NBLAST search with Cha-F-600134 as query (black). Neuron plots of (from higher to lower score): the query and top hit, top 8 hits, hits with a score over 5,000 and hits with a score over 0. The top hit corresponds to an image of a neuron from the same brain but from a different raw image. It is very similar to the query neuron. As the score decreases, so does the similarity of the hits to the query. On the right, histogram of forward scores. Only hits with scores over -8,000 are shown. The score of the query and top hit are indicated. A dashed purple line marks 0. The left inset shows a zoomed view of the top hits (score > 5,000) (dashed blue rectangle in main plot). The score of the query, top hit and second top hits are indicated.

**Figure S2:**
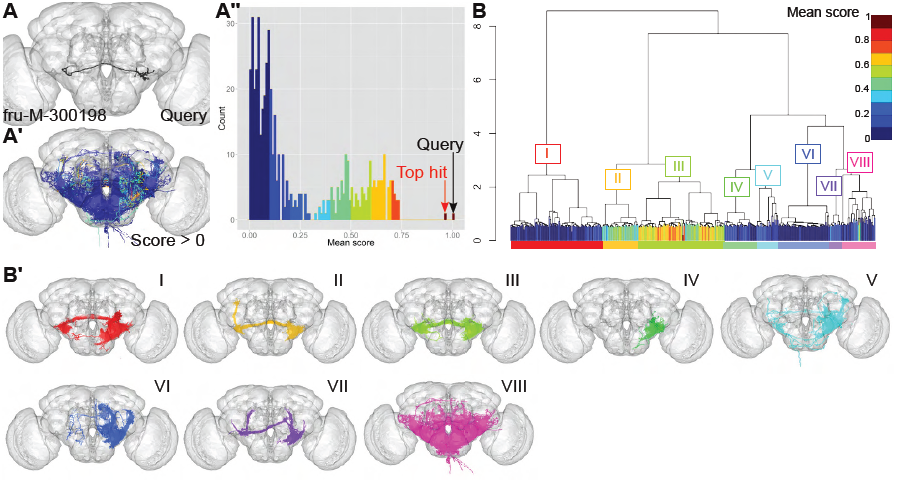
NBLAST search using mean scores, related to Figure 3. **(A)** The query neuron fru-M-300198 (same as in Figure 3B). **(A’)** Neuron plot of the hits with a mean score over 0 for a search against the query. Hits are colored by score bin (10), as in A”. **(A”)** Histogram of mean scores for hits against fru-M-300198 with a score over 0 divided into 10 bins (indicated in the scale bar in B). **(B)** Hierarchical clustering of hits with a mean score over 0. The leaves of the dendrogram are colored by score (same as in A”), and as shown in the scale bar. The dendrogram was divided into eight groups (I–VIII), with each one being assigned a color, shown on the colored rectangle below the leaves. The query neuron is in group III, and the hits with the higher scores are in groups II and III. **(B’)** Neuron plots corresponding to the dendrogram groups (I–VIII), following the colors assigned to each group. Groups II and III, corresponding to the highest scores, are the most similar neurons to the query.

**Figure S3:**
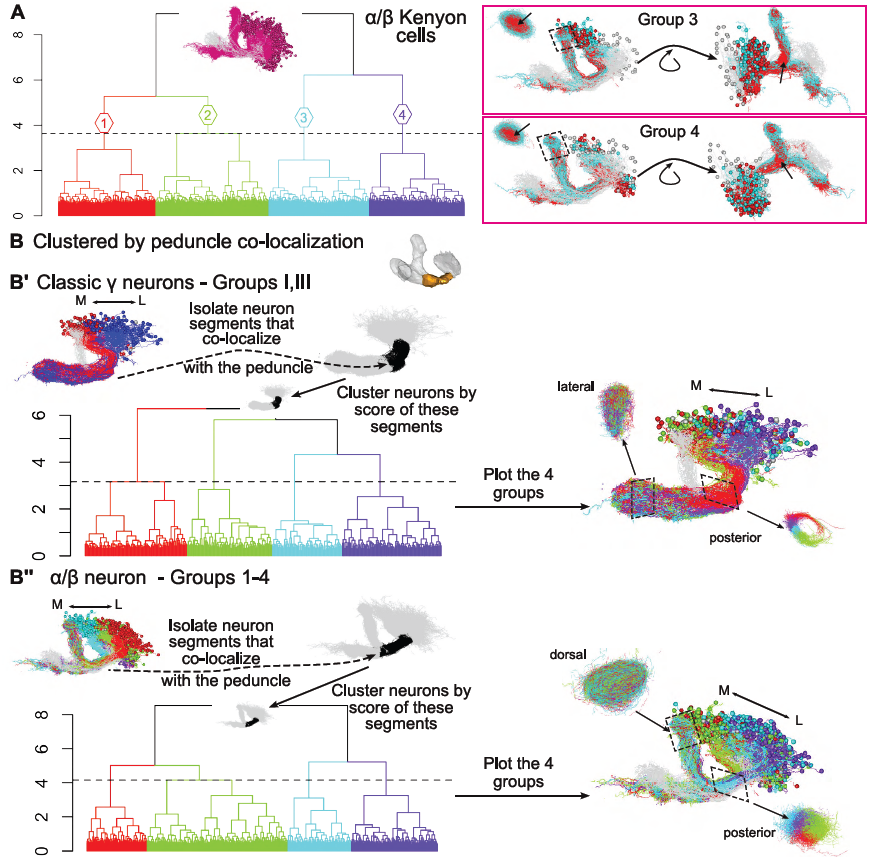
NBLAST search and classification of hits reveals Kenyon cells subtypes, related to Figure 4. **(A)** Hierarchical clustering (HC) of the α/β neurons, divided into four groups (1–4) (h=3.64) (same as in Figure 4D’). The inset on the dendrogram shows the α/β neurons. Groups 3 and 4 were clustered and divided into 2 groups each. This separates the neurons into peripheral (cyan) and core (red) in the α lobe. Peripheral neurons occupied a more lateral calyx position and were dorsal to core neurons in the peduncle and β lobe. Similar analysis to group 1 is shown in Figure 4D”. Lateral oblique, posterior oblique and a dorsal view of a peduncle slice (position indicated by dashed rectangle) are shown. **(B)** Reclustering of Kenyon cells based on the co-localization of neurons segments in the peduncle. The neuron segments that co-localize with the peduncle were isolated, followed by HC of the neurons based on the NBLAST score of the segments. **(B’)** HC of the neurons segments of classic γ neurons, groups I and III (see Figure 4B”) divided into 4 groups, h=3.16. Neuron plot of the 4 groups. A posterior view of a slice of the peduncle shows an expected clear organization. It correlates to the position of the neurons in the calyx, with more medial neurons (cyan and green) being dorsal and ventral in the peduncle than more lateral neurons (red and purple). No clear structural organization is discernible in a lateral view of a slice of the γ lobe. **(B”)** HC of the neurons segments of classic α/β neurons, groups 1 to 4 (see Figure 4D’) divided into 4 groups, h=4.16. Neuron plot of the 4 groups. A posterior view of a peduncle slice shows an expected clear organization. It correlates to the position of the neurons in the calyx, with more medial neurons (cyan and green) being ventrolateral in the peduncle than more lateral neurons (red and purple). No structural organization is discernible in a dorsal view of a slice of the α lobe. For all neuron plots, the neurons in grey correspond to the Kenyon cell exemplars.

**Table S1:** Correspondences between hierarchical clustering groups of AOTU- and PVLP-innervating (groups 1-7) and PVLP- and PLP-innervating (groups A-G) uVPNs via NBLAST scores and previously determined neuron types (Otsuna and Ito, 2006). Related to Figure 6 and Figure S4.

| Dendrogram group | Neuron type | Comments |
| --- | --- | --- |
| 1 | LC6 | Lobula innervation, dorsal cell bodies, axons follow the anterior optic tract to the AOTU, turning ventrally midway to innervate the lateral PVLP. |
| 2 | LC9 | As group 1, but terminating in the medial PVLP rather than extending laterally. |
| 3 | LC10B (possibly different subtype to 4) | Dorsal AOTU innervation, ventral to group 4. |
| 4 | LC10B (possibly different subtype to 3) | Dorsal AOTU innervation, dorsal to group 3. |
| 5 | New LC10 subtype | The most ventral AOTU innervation. Similar to group 7, but ventral to it in the AOTU. |
| 6 | LC10A | Axons project through ventral AOTU, turn sharply dorsally to terminate in the dorsal AOTU. |
| 7 | New LC10 subtype | Ventral AOTU innervation, dorsal to group 5. |
| A | LC12 subtype | Innervation in most lateral and anterior PVLP glomerulus. Note: corresponds to LC17 in Panser et al. (2015). |
| B | LC12 subtype | Innervation in more posterior and medial PVLP glomerulus than group A. Note: corresponds to LC12 in Panser et al. (2015). |
| C | LC4 | Innervates a more medial PVLP glomerulus than LC12. |
| D | LT12 | Tentative match. Class was identified in Otsuna and Ito (2006) based on a single neuron. |
| E | New LC class | Innervates more dorsal PVLP glomerulus than LC12. Extends along the posterior PVLP, with a sharp anterior turn. Terminates with a blunt-stick like ending in the lateral PVLP. Corresponds to LC18 in Panser et al. (2015). |
| F | New LT class | Similar to LT12, but with projections posterior to it, terminating in the lateral region of the superior posterior slope. |
| G | Unmatched | Do not correspond to a single type. |

**Figure S4:**
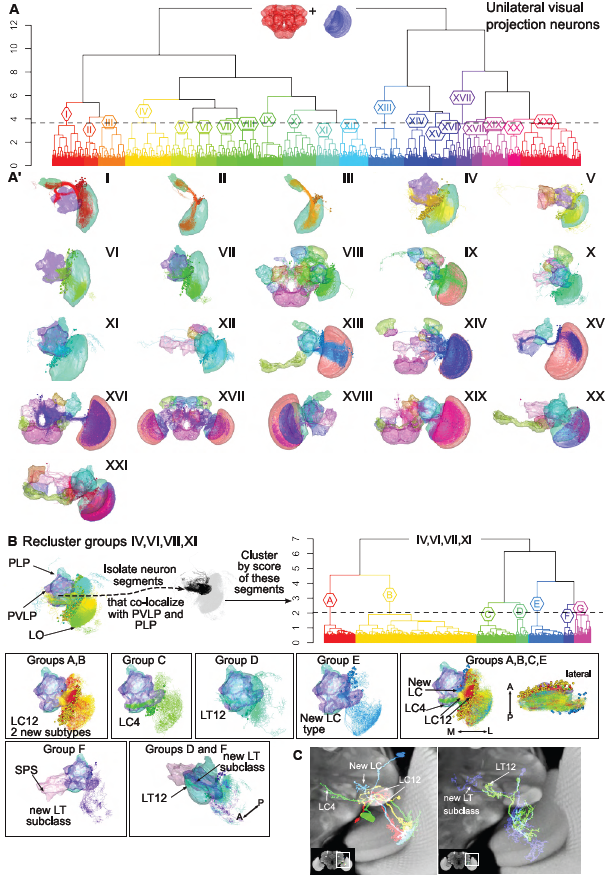
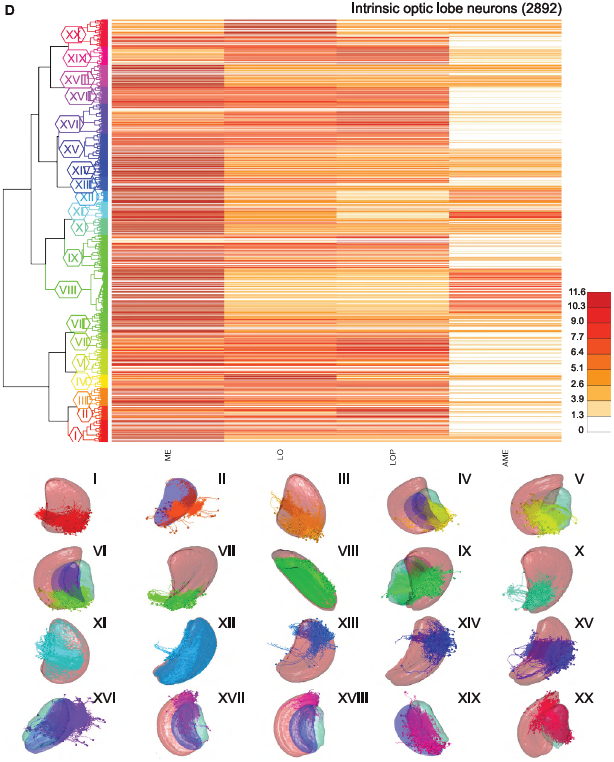
NBLAST search and classification hits uncovers unilateral visual projection neurons neuronal types, related to Figure 6. **(A)** Hierarchical clustering (HC) analysis of unilateral visual projection neurons (uVPNs). The dendrogram was divided into 21 groups (I–XXI), h=3.65. Inset on the dendrogram shows the neuropils considered for the overlap. Below, neuron plots of groups I to XXI. The neuropils that contain the most overlap are shown. **(B)** Reclustering of uVPN groups IV, VI, VII and XI from A. The neuron segments that colocalize with either the PVLP or PLP were isolated, followed by HC of the neurons based on the NBLAST score of these neuron segments. The dendrogram was divided into seven groups (A–G), h=2.04. Neuron plots matching the dendrogram groups to known uVPN types. An anterior, a lateral or lateral oblique views are shown. Groups A and B possibly correspond to two LC12 subtypes. Group B innervates a more anterior and medial glomeruli than group A (see also C). Group C corresponds to LC4. Group D corresponds to LT12 neurons. Group E corresponds to a new LC type. Group F corresponds to a possibly new LT subclass. **(C)** Overlay of Z projections of registered image stacks of example neurons from the types identified in B on a partial Z projection of the template brain (a different one for each panel). The white rectangle on the inset shows the location of the zoomed in area. LC: lobula columnar neuron; LT: lobula tangential neuron. **(D)** Hierarchical clustering of intrinsic optic lobe neurons. This neuron set was defined as any neuron that overlapped only one of the optic lobes and with no arborization in the central brain neuropils. Dendrogram of the intrinsic optic lobe neurons, divided into into 20 groups (I–XX) with the corresponding heatmap calculated from the neuropil overlap in the different neuropils: medulla (ME), lobula (LO), lobula plate (LOP) and accessory medulla (AME). Neuron plots corresponding to the dendrogram groups are shown below. The neuropils for which the overlap is more significant are plotted. Although some organizational structure is seen, the dendrogram groups to do not represent unique types.

**Figure S5:**
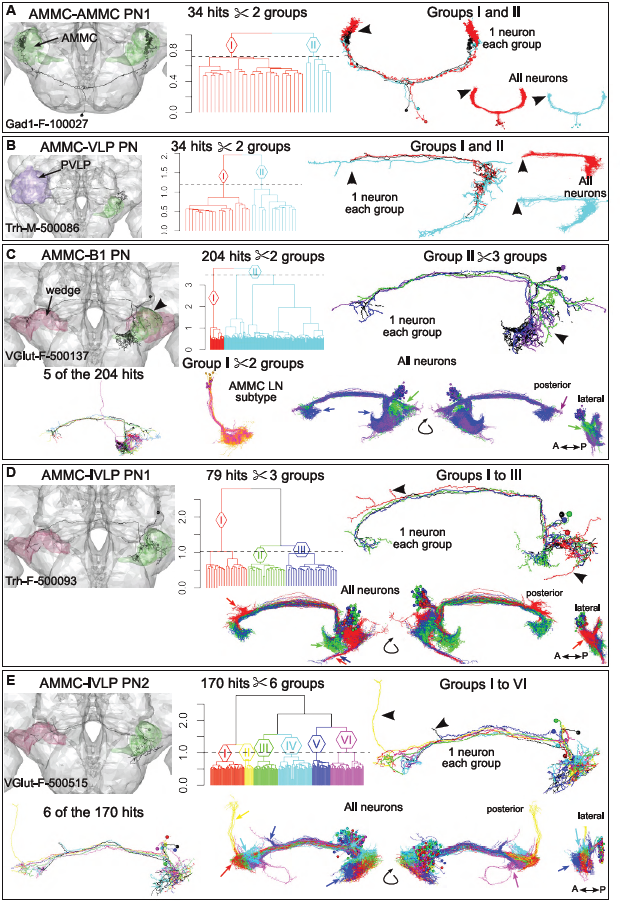
NBLAST search and classification hits reveals auditory neuron types, related to Figure 6. On the left, a neuron of the types identified by Lai et al. (2012) (A-E) which was used as query. The dendrogram and neuron plots for each type (anterior, posterior and lateral views) showing either one or all neurons from each clustering group are shown on the middle and right panels. Main differences in arborization between neurons are indicated by arrowheads and arrows. **(A)** Hierarchical clustering (HC) of AMMC-AMMC projection neuron 1 (PN1). The 34 top scorers were clustered and divided into 2 groups, h=0.72. Neuron plot of the query neuron (black), and one neuron from each group (red and cyan). To the right, neuron plots of of each group. **(B)** HC of AMMC-VLP PNs. The 34 top scorers were clustered and divided into 2 groups, h=1.2. Neuron plot of the query neuron (black), and one neuron from each group (red and cyan). On the left, neuron plots of each group. **(C)** HC of AMMC-B1 PNs. Five hits of the 204 top scorers are shown on the left. The 204 top scorers were clustered and divided into 2 groups, h=3.46. Group I was matched to an unidentified type of AMMC local neurons (LN). It was clustered and divided into 2 groups, h=0.77. Group II corresponds to a mix of AMMC-B1 PNs and AMMC-IVLP PN1. After selecting the AMMC-B1 PNs, the neurons were clustered and divided into 3 groups, h=1.5. Neuron plot of the query neuron (black), and one neuron from each group (purple and green). Below, neuron plots of the 3 groups. **(D)** HC of AMMC-IVLP PN1. The 79 top scorers were clustered and divided into 3 groups, h=1.03. Neuron plot of the query neuron (black), and one neuron from each group (red, green and blue). Below, anterior, posterior and lateral view neuron plots of the 3 groups. **(E)** HC of AMMC-IVLP PN2. Six hits of the 170 top scorers are shown on the left. The 170 top scorers were clustered and divided into 6 groups, h=1.02. Neuron plot of the query neuron (black), and one neuron from each group (red, yellow, green, cyan, blue, magenta). Below, anterior, posterior and lateral view neuron plots of the 6 groups. Arrows and arrowheads indicate differences between groups. wedge in magenta, AMMC in green or PVLP in purple

**Table S2:**
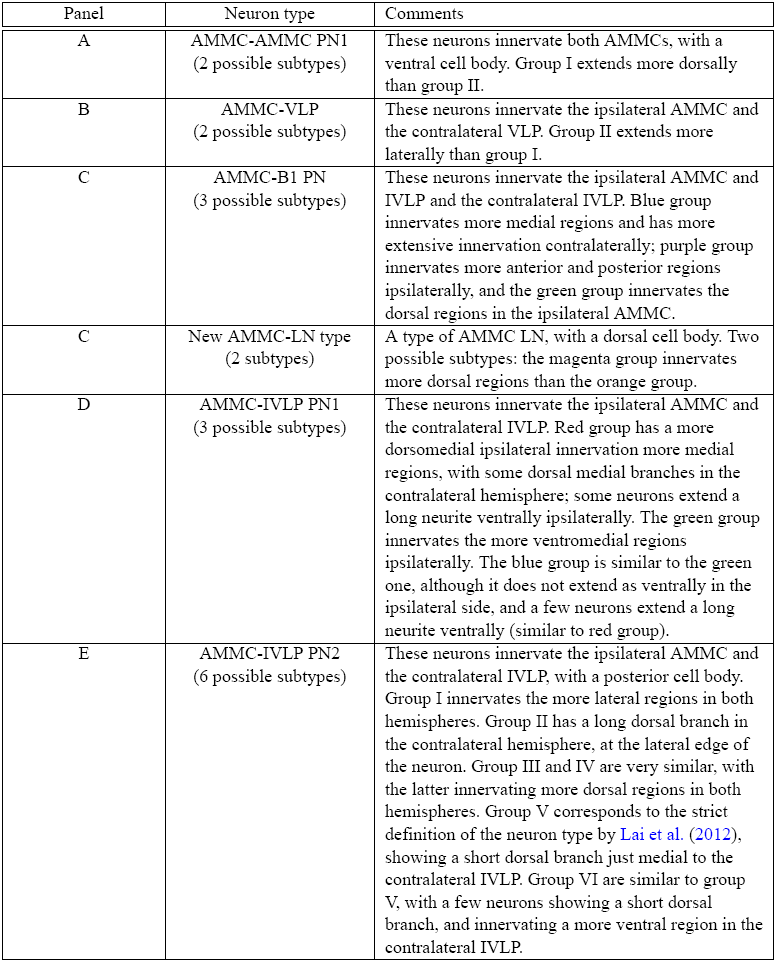
Correspondence between hierarchical clustering of auditory neuron via NBLAST scores and previously determined neuron types (Lai et al., 2012). Related to Figure S5.

